# Medea: An AI agent for therapeutic reasoning across biological contexts

**DOI:** 10.64898/2026.01.16.696667

**Authors:** Pengwei Sui, Michelle M. Li, Brenton P. Munson, Shanghua Gao, Wanxiang Shen, Valentina Giunchiglia, Andrew Shen, Yepeng Huang, Zhenglun Kong, Katherine Licon, Trey Ideker, Marinka Zitnik

## Abstract

Therapeutic hypotheses can transfer across diseases but their relevance depends on biological context. The same target, perturbation, or treatment can produce different effects across cell types, disease states, genetic backgrounds, and patients. Therapeutic reasoning therefore requires methods that preserve context, test when evidence supports transfer, and identify where context-specific effects limit it. Although AI agents can perform therapeutic analyses, existing systems often fail to preserve biological context over long workflows, verify intermediate computational steps, or reconcile conflicting evidence across datasets and literature. Here, we present Medea, an AI agent for therapeutic reasoning across biological contexts. Medea executes multi-step analyses using biological tools, machine learning models, and literature retrieval while enforcing verification during planning, execution, and evidence synthesis. We evaluate Medea across 5,673 open-ended analyses in three domains: cell type specific therapeutic target nomination in five diseases and 29 cell types, synthetic lethality prediction in 7 cancer cell lines, and immunotherapy response prediction from multimodal patient profiles. Using a previously unpublished epistatic miniarray profiling screen performed under two DNA-damaging treatments, we evaluate Medea on predicting synthetic lethality among 238,046 gene-gene pairs in yeast. Medea predicts these experimentally measured synthetic lethal interactions, indicating that its performance reflects biological relevance rather than information leakage from benchmark datasets. Across these evaluations, Medea improves performance over large language models, reasoning models, biomedical agents, and specialized machine learning models while maintaining low failure rates and calibrated abstention. These results show that verifiable AI agents can perform therapeutic analyses across biological contexts.

## Introduction

Therapeutic reasoning requires assessing evidence across biological contexts [1–3]. A drug target or therapeutic perturbation that is effective in one context may show limited efficacy or unacceptable safety risks in another, depending on disease state, tissue environment, and patient medical history. For example, response to PD-1 or PD-L1 blockade can vary substantially across patients with similar tumor mutational burden, because treatment response also depends on tumor microenvironment programs, including interferon signaling, T-cell exhaustion, and immune cell infiltration [4–6]. Similarly, perturbing the same gene can produce distinct effects across cell types [2, 3, 7]. These context-dependent effects determine which candidate targets or perturbations should be prioritized, in which cellular models or patient populations, and under which molecular conditions. Omics datasets provide the empirical foundation for therapeutic reasoning by linking molecular readouts to cellular phenotypes, disease mechanisms, and patient responses [8–13]. AI agents are beginning to help explore omics datasets by translating therapeutic hypotheses into research plans, retrieving relevant data, and executing analyses [14–22]. However, many agents still rely on large language model (LLM) parametric memory rather than grounding intermediate results in omics datasets, or they follow fixed computational biology workflows [23].

Three gaps limit how these agents can support therapeutic reasoning from omics data. First, agents often lose track of biological context over long-horizon analyses. They do not preserve the specified cell type, disease phenotype or patient characteristics across agentic workflow, and may therefore apply single-cell tools to bulk questions or query pathway knowledge bases curated for unrelated tissues. These context shifts can bias analyses toward marker genes from abundant cell populations rather than targets in disease-relevant cell types [18, 20]. Second, agents rarely verify feasibility before and after execution. Pre-run checks often do not validate tool-dataset assumptions, parameter compatibility, or statistical requirements, whereas post-run checks frequently stop at catching runtime failures. As a result, an analysis can execute successfully while still producing biologically invalid conclusions, for example when differential expression analyses use mismatched covariates that propagate into downstream pathway ranking [21, 22]. Third, agents can struggle to reconcile evidence across datasets, tools, and literature sources, often aggregating studies without screening for contextual relevance or experimental evidence [18, 19].

Here, we present Medea, an AI agent for therapeutic reasoning across biological contexts (Figure 1a). Medea takes an omics objective and executes a multi-step analysis with verification throughout planning, execution, and evidence synthesis. Medea comprises four modules: ResearchPlanning, which specifies biological context and verifies plan integrity; Analysis, which executes tool-grounded analyses with pre-run and post-run validation; LiteratureReasoning, which retrieves and screens studies for contextual relevance; and MultiRoundDiscussion, which reconciles evidence across tool outputs, literature, and language model reasoning or abstains when evidence is insufficient. Medea operates over a tool space spanning single-cell and bulk transcriptomics, protein interaction networks, pathway and ontology knowledge bases, cancer dependency maps, and foundation models, and we evaluate it across three open-ended domains.

**Figure 1:**
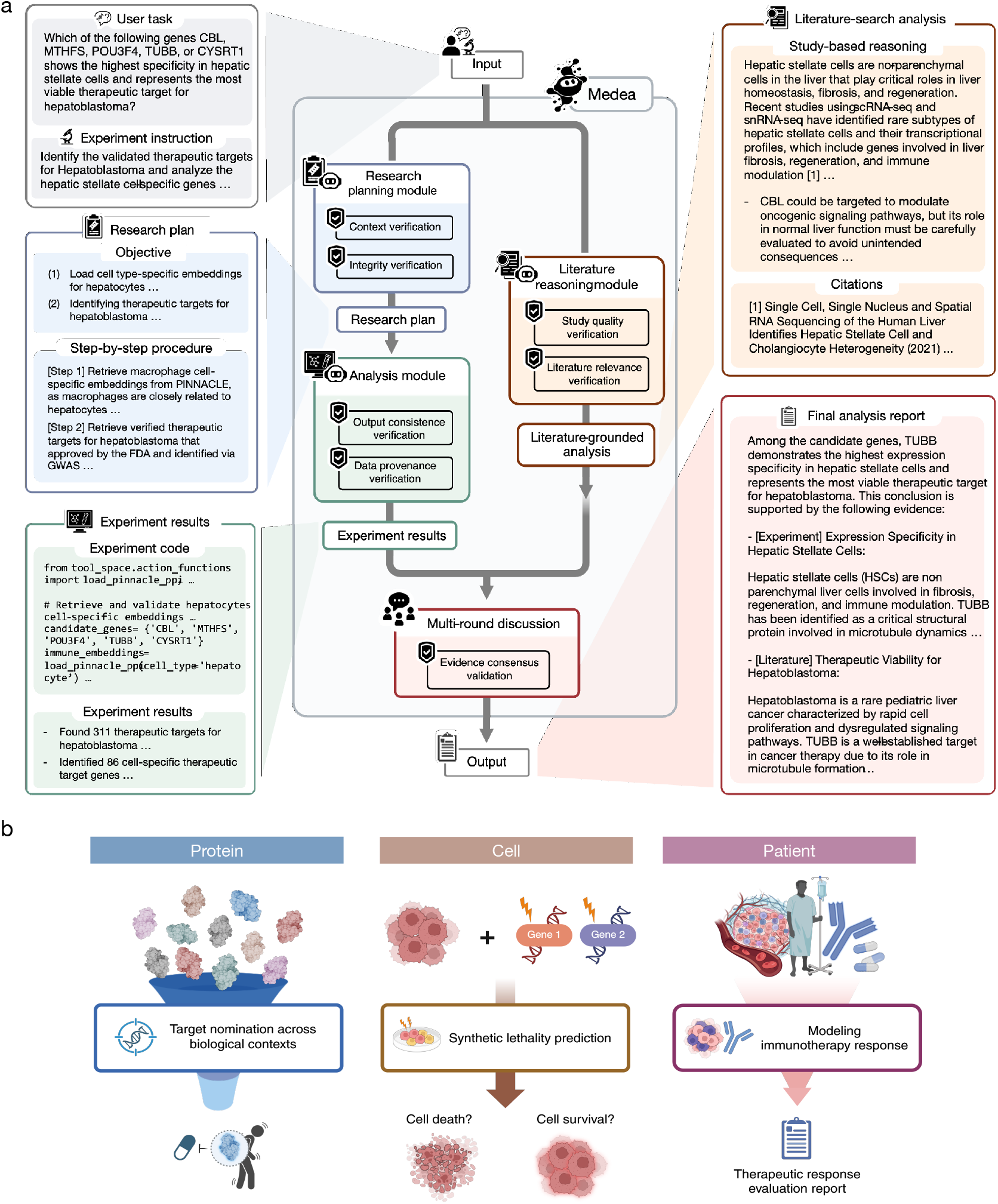
Overview of Medea and open-ended evaluation domains. **(a)** Medea takes an omics objective and an optional experiment instruction, produces a research plan, executes omics analyses using tools, retrieves and screens literature, and reconciles evidence to return a final report or calibrated abstention. Medea consists of four modules: ResearchPlanning (context and integrity verification for plan construction), Analysis (tool execution with pre-run checks and post-run verification), LiteratureReasoning (literature retrieval with relevance and evidence-strength assessment), and MultiRoundDiscussion (evidence reconciliation across module outputs). **(b)** Medea is evaluated on three domains: cell type specific target nomination, synthetic lethality reasoning in cell lines, and immunotherapy response prediction from patient tumor transcriptomes and clinical data.

In cell type specific target nomination, Medea prioritizes candidate targets in disease-relevant cell types rather than tissue-level averages. Across 2,400 analyses covering five diseases and primary cell types, Medea improves the accuracy of LLMs by up to 45.9% in rheumatoid arthritis and 33.0% in Sjögren’s Syndrome. In synthetic lethality prediction, Medea integrates genetic interaction signals with pathway evidence to identify gene pairs whose combined inhibition is predicted to impair cancer cell viability. In 2,379 analyses in seven cell lines, Medea improves accuracy by up to 20.2% in MCF7 and 13.9% in A549, with lower failure rates than the LLM alone. In immunotherapy response prediction, Medea links tumor-intrinsic and microenvironment programs to treatment responses by analyzing immunological programs related to antigen presentation, interferon signaling, and T-cell exhaustion. Across 894 analyses involving 298 patients, Medea achieves up to 23.9% higher accuracy than existing models.

We then leverage Medea for predicting synthetic lethality among 238,046 gene-gene pairs in yeast, whose genetic interaction maps have been used to reconstruct the wiring of DNA repair and other core pathways [24]. Bleomycin and dimethyl sulfate induce double-strand breaks and DNA alkylation damage so interactions specific to these treatments point to genes with redundant or buffering roles in the corresponding repair pathways rather than in viability generally [25]. We measure these interactions using a previously unpublished epistatic miniarray profile screen performed under bleomycin and dimethyl sulfate exposure, against which we evaluate Medea’s predictions. In non-abstention cases, Medea outperforms GPT-5 and GPT-5.2 in predicting these experimentally measured synthetic lethal interactions, showing that Medea’s performance reflects biological relevance rather than information leakage from benchmark datasets.

Ablation analyses show complementary contributions from Medea’s agentic modules. A literature-only configuration of Medea abstains in 79.1% of disease contexts, whereas an LLM-only configuration abstains in 1.8% of analyses yet accounts for the largest share of errors. The full configuration of Medea achieves the best performance with the lowest failure rate.

## Results

### Medea agent for therapeutic reasoning across biological contexts

Medea takes as input an omics-based objective specified as a natural language instruction and an optional experiment instruction. To support different levels of user expertise, Medea can accept either a high-level objective alone or an experiment instruction that specifies datasets, tools, analytical steps, or constraints to follow during execution. When no experiment instruction is provided, Medea generates and verifies its own research plan before executing the analysis. Medea couples agentic modules with tool use (Figure 1a) and can invoke any of 27 tools (Methods 1.1, Figure 2a) to execute multi-step analyses, returning a report grounded in outputs from machine learning models [26–28], omics datasets [29–38], and literature [39–41].

**Figure 2:**
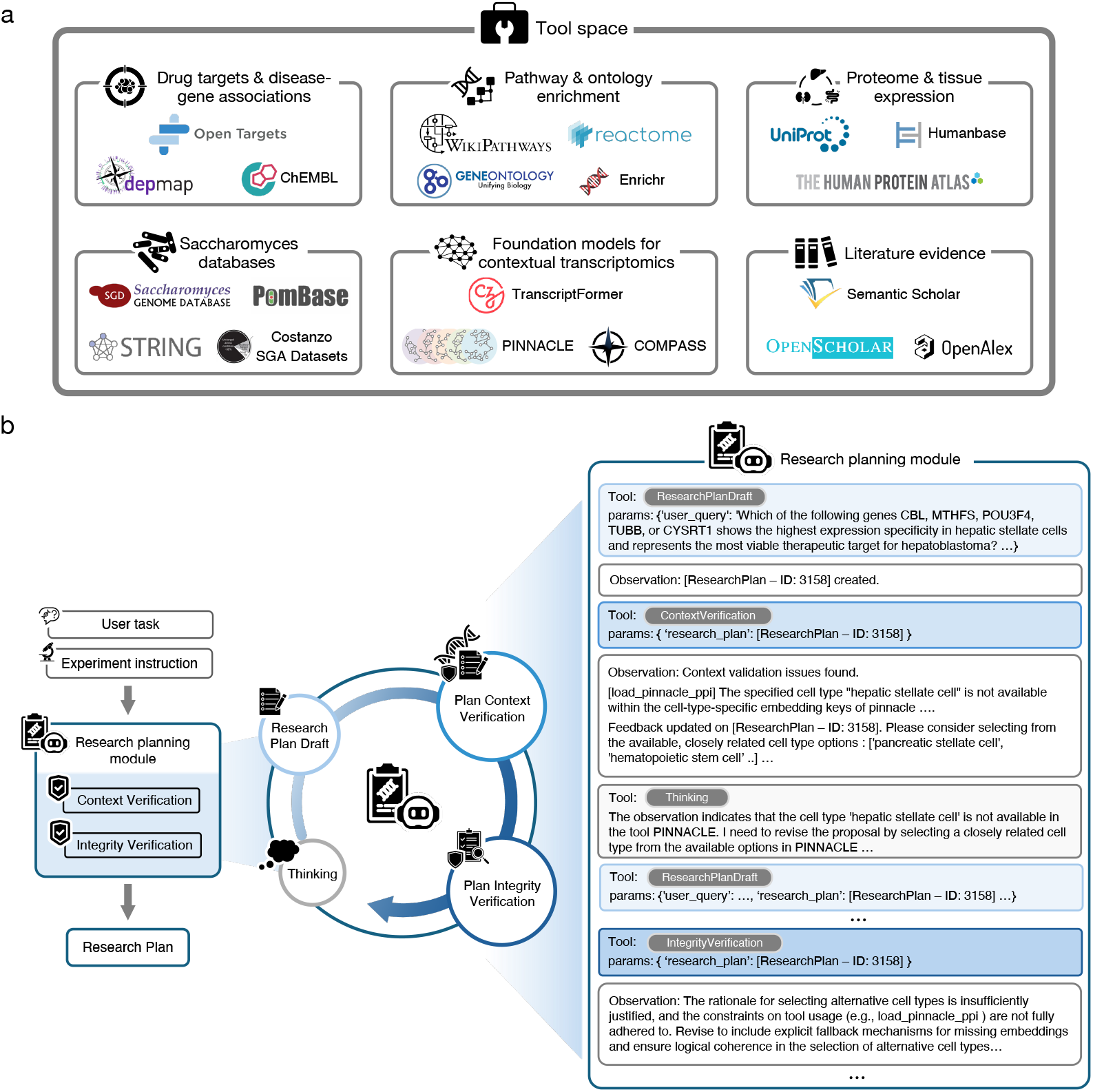
Tool space and verified planning in Medea. **(a)** Medea uses a global tool space (Methods 1.1) with resources about therapeutic targets, disease associations, proteomics, and tissue expression; tools to perform gene set enrichment and pathway analyses; machine learning models for single-cell and bulk omics; and tools for literature retrieval. **(b)** Given an omics objective (user instruction) and an optional experiment instruction, the ResearchPlanning module (Methods 1.2) generates a multi-step analysis plan. Context verification checks tool and data compatibility with the research objective. Integrity verification audits the research plan’s feasibility, completeness, and logical consistency.

Medea orchestrates four modules: ResearchPlanning (Methods 1.2), Analysis (Methods 1.3), LiteratureReasoning (Methods 1.4), and MultiRoundDiscussion (Methods 1.5). The ResearchPlanning module transforms an omics-based objective into an executable computational experiment plan. It defines the biological context, breaks the objective into subtasks, selects tools for each step (e.g., databases, machine learning models), and checks whether the plan is specific, feasible, and logically valid [42–45]. The Analysis module translates the plan into tool-calling analyses that use databases, APIs, machine learning models, and other agents. It checks code and tool dependencies before execution (syntax and dependency checks), runs analyses in a sandboxed environment with error capture, and performs post-execution verification (data provenance auditing [46]) to ensure outputs remain aligned with the plan and the omics objective. The LiteratureReasoning module retrieves and screens literature using Semantic Scholar [39] and OpenAlex [40], then uses OpenScholar [41] to synthesize evidence from relevant studies. Finally, Medea queries the backbone LLM to obtain a parametric-knowledge response and passes this response, the Analysis output, and the LiteratureReasoning output to MultiRoundDiscussion. MultiRoundDiscussion runs a multi-round deliberation process [47] in which a panel of LLMs evaluates the evidence streams, identifies agreement and conflict, and produces a consensus answer. When the evidence streams do not support a single answer, for example because tool outputs are weak, literature evidence is missing or conflicting, or the panel cannot reach agreement, MultiRoundDiscussion returns an abstention rather than forcing a conclusion. In evaluations, Medea can activate any subset of modules and tools to complete omics objectives, a capability we evaluate across three open-ended domains: cell type specific target nomination, synthetic lethality reasoning, and immunotherapy response prediction.

### Drug target nomination in disease-relevant cell types

Nominating drug targets requires reasoning about the disease and cellular context in which a perturbation is expected to act. The same target can produce different therapeutic or toxic effects across tissues, cell types, and cellular states [1, 48–50]. Yet many computational approaches for target prediction rely on bulk tissue measurements or established cell lines and therefore lack the resolution needed to evaluate therapeutic hypotheses at the level of specific cell types [7, 51, 52]. We therefore evaluate whether AI agents can complete omics objectives that require reasoning about candidate therapeutic targets within disease-relevant cellular contexts (Figure 3a).

**Figure 3:**
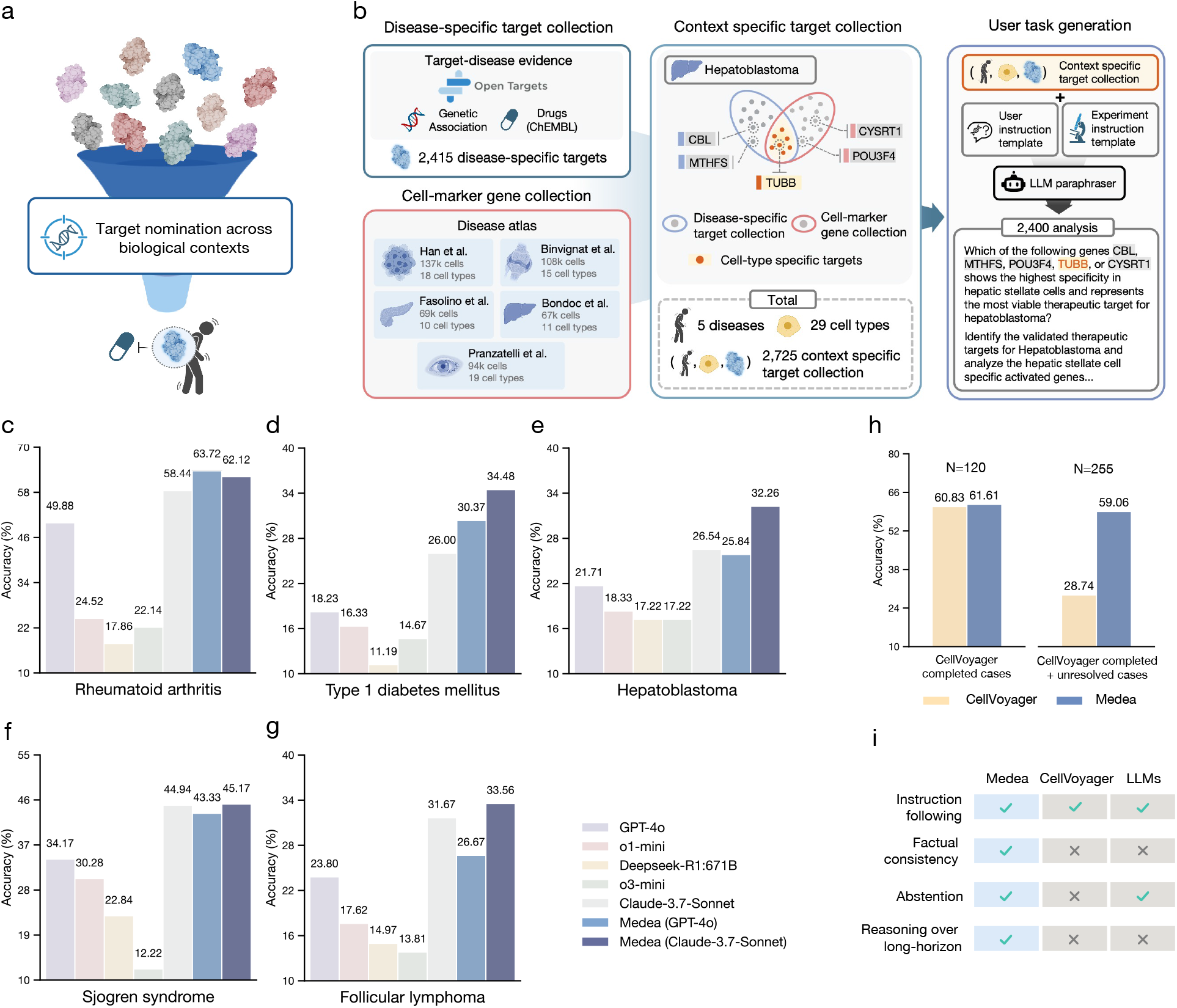
Cell type specific target nomination benchmark. **(a)** Given a disease, a cell type, and a set of candidate genes, Medea executes the omics objective of nominating the most supported target for the specified context by integrating evidence from databases, single-cell foundation models, and the scientific literature (Methods 2). **(b)** Constructing the benchmark dataset of context-specific targets across five diseases and multiple primary cell types leverages single-cell atlases and resources about therapeutic targets and disease-gene associations. **(c-h)** Performance of Medea compared to five LLMs and a single-cell computational biology agent, CellVoyager [20], across disease and cell type contexts. **(i)** Qualitative comparison of capabilities relevant to this benchmark, including instruction following, factual consistency checks, calibrated abstention, and long-horizon multi-step reasoning.

We construct an evaluation benchmark spanning 29 cell types across five diseases: rheumatoid arthritis (RA) [53], type 1 diabetes mellitus (T1DM) [54], Sjögren’s syndrome (SS) [55], hepatoblastoma (HB) [56], and follicular lymphoma (FL) [57] (Figure 3b; Methods 2). For each disease, we retrieve single-cell transcriptomic atlases from healthy individuals and patients and identify differentially expressed genes within each cell type between healthy and disease states. We then construct context-specific candidate targets by combining differential expression with prior disease and therapeutic evidence from human genetics [58] and ChEMBL [59]. For each disease–cell-type context, we generate 60 analyses in which the agent receives a disease, a cell type, and five candidate genes, consisting of one candidate target supported by differential expression and prior therapeutic evidence, and four negative candidates. This process yields 2,400 analyses in total: 420 across 7 cell types for RA, 600 across 9 cell types for T1DM, 360 across 6 cell types for SS, 180 across 3 cell types for HB, and 840 across 14 cell types for FL. In each analysis, the model or agent selects the gene with the strongest evidence of therapeutic potential in the specified disease and cell type and provides a rationale. We evaluate performance using per-analysis accuracy, defined as the fraction of analyses in which the evidence-supported target candidate is nominated among the five candidates. For agents that can abstain when confidence is low, we additionally report abstention rates and accuracy conditioned on non-abstention.

Medea is evaluated against five LLMs and a computational biology agent, CellVoyager [20], for target nomination in disease-relevant cell types. For each of the 2,400 analyses, Medea generates a target nomination together with a report that summarizes the supporting evidence in the specified disease and cell type context.

Medea outperforms LLMs, reasoning models, and CellVoyager on cell type specific target nomination (Figure 3; Methods 5.2). Medea improves accuracy over LLMs by up to 45.9% in rheumatoid arthritis (p-value *<* 0.0001 using McNemar’s test [60]; Figure 3c), 23.3% in type 1 diabetes mellitus (p-value *<* 0.0001; Figure 3d), 15.0% in hepatoblastoma (p-value *<* 0.0001; Figure 3e), 33.0% in Sjögren’s Syndrome (p-value *<* 0.0001; Figure 3f), and 19.8% in follicular lymphoma (p-value *<* 0.0001; Figure 3g). We additionally evaluate Medea with GPT-4o or Claude 3.7 Sonnet as the backbone LLM. Medea outperforms its backbone LLM, which is defined as the underlying language model used on its own without the ResearchPlanning, Analysis, or LiteratureReasoning modules. For example, Medea (GPT-4o) and Medea (Claude 3.7 Sonnet) are 12.1% (*p*-value *<* 0.0001) and 8.5% (*p*-value = 0.0004) more accurate than GPT-4o and Claude 3.7 Sonnet alone, respectively, on the omics objectives for type 1 diabetes mellitus target nomination (Figure 3d). These results show that integrating verification-aware modules and tools improves target nomination across disease and cell type contexts.

We also evaluate CellVoyager [20] for target nomination for rheumatoid arthritis [26]. We compare CellVoyager against Medea (Claude 3.7 Sonnet) using the global tool space (Section 1.1). Because CellVoyager produces lengthy multimodal outputs and exhibits high failure rates, a human expert reviews and scores all CellVoyager outputs. Across 420 analyses, CellVoyager completes 28.6% (120 out of 420). On these 120 completed analyses, CellVoyager is 0.8% less accurate than Medea (Figure 3h). In 39.3% of analyses (165 out of 420), CellVoyager fails to nominate any target due to errors during data preprocessing; on the remaining 255 analyses, Medea is 30.3% (or 2.1 times) more accurate than CellVoyager (Figure 3h). We also find that CellVoyager does not enforce factual consistency [61–63] (Figure 3i; Supplementary Note 6); for example, it often hallucinates gene names. Finally, while LLMs can be instructed to abstain when uncertain, CellVoyager is not designed to abstain [64–66] (Figure 3i), and struggles with long-horizon reasoning [42, 67, 68] because it only interprets the output of the most recent Jupyter cell (Figure 3i). In contrast, Medea maintains factual consistency at each step of the analysis, can abstain when evidence is insufficient, and completes analyses that require multi-step reasoning over long horizons, which correspond to higher accuracy and lower failure rates on the benchmark.

### Context verification improves cell type level performance

We assess Medea’s ability to reason about candidate targets within the user-specified cell type context, which is critical for therapeutic efficacy and safety. Completing this omics-based objective requires correctly identifying the stated context and retrieving context-appropriate evidence [18]. Closely related cell types can have distinct roles; for example, naïve CD4^+^ *αβ* T cells versus effector memory CD4^+^ *αβ* T cells and naïve CD8^+^ *αβ* T cells [69, 70]. However, LLMs often miss such distinctions [18, 23] and may default to higher-level lineages, such as CD4^+^ or CD8^+^ T cells. Medea is designed to perform context verification [71, 72] so that intermediate decisions remain consistent with the specified cell type and disease context. To quantify the contribution of context verification, we stratify Medea’s performance across the cell type and disease contexts in the target nomination benchmark (Methods 2).

Medea performs comparably or better than GPT-4o in diverse cell type and disease contexts (Figure 4a). Medea boosts the accuracy of predicting therapeutic targets for rheumatoid arthritis in myeloid dendritic cells by 28.9%, naïve CD4^+^ *αβ* T cells by 21.7%, effector memory CD4^+^ *αβ* T cells by 21.1%, and naïve CD8^+^ *αβ* T cells by 12.2%. Performance gains in nominating therapeutic targets for rheumatoid arthritis within these cell types are meaningful. The localization of certain myeloid dendritic cells in the synovium of patients with rheumatoid arthritis has been associated with immune homeostasis [73]. As different subsets of CD4+ [74] and CD8^+^ T cells [75] contribute uniquely to the pathogenesis of rheumatoid arthritis, it is important to consider granular subtypes (e.g., naïve versus effector memory CD4^+^ *αβ* T cells) for nominating therapeutic targets. Medea and GPT-4o perform comparably on naïve B cells and natural killer (NK) cells. For type 1 diabetes mellitus, Medea consistently outperforms GPT-4o on all nine cell type contexts, with performance gains of up to 21.7%. For Sjögren’s Syndrome, Medea yields improved average accuracies by 29.6% in endothelial cells, 15% in fibroblasts, and 14.8% in IgA plasma cells. Endothelial cells and fibroblasts contribute to the recruitment of or interact with lymphocytes in salivary glands, which are particularly affected by Sjögren’s Syndrome, respectively [76]. Since IgA plasma cells are enriched in patients with Sjögren’s Syndrome, they serve as one of the histopathological features for diagnosis [77, 78] and a potential target for treating systemic autoimmune rheumatic diseases [79]. Medea’s accuracy doubles (7.0% vs. 13.6%) when nominating therapeutic targets for hepatoblastoma in Kupffer cells, which are the liver’s first line of defense [80]. For follicular lymphoma, Medea achieves 15.0% and 13.9% higher average accuracy in myeloid cells and plasmacytoid dendritic cells than GPT-4o, respectively. Myeloid cells are manipulated by tumor cells to promote tumor angiogenesis, cell invasion, and metastasis [81], and the activation of plasmacytoid dendritic cells can boost innate and adaptive cancer immunity [82]. These findings indicate that Medea identifies the disease-relevant cell type context, retrieves and executes context-appropriate tools, and reasons within the specified biological context.

**Figure 4:**
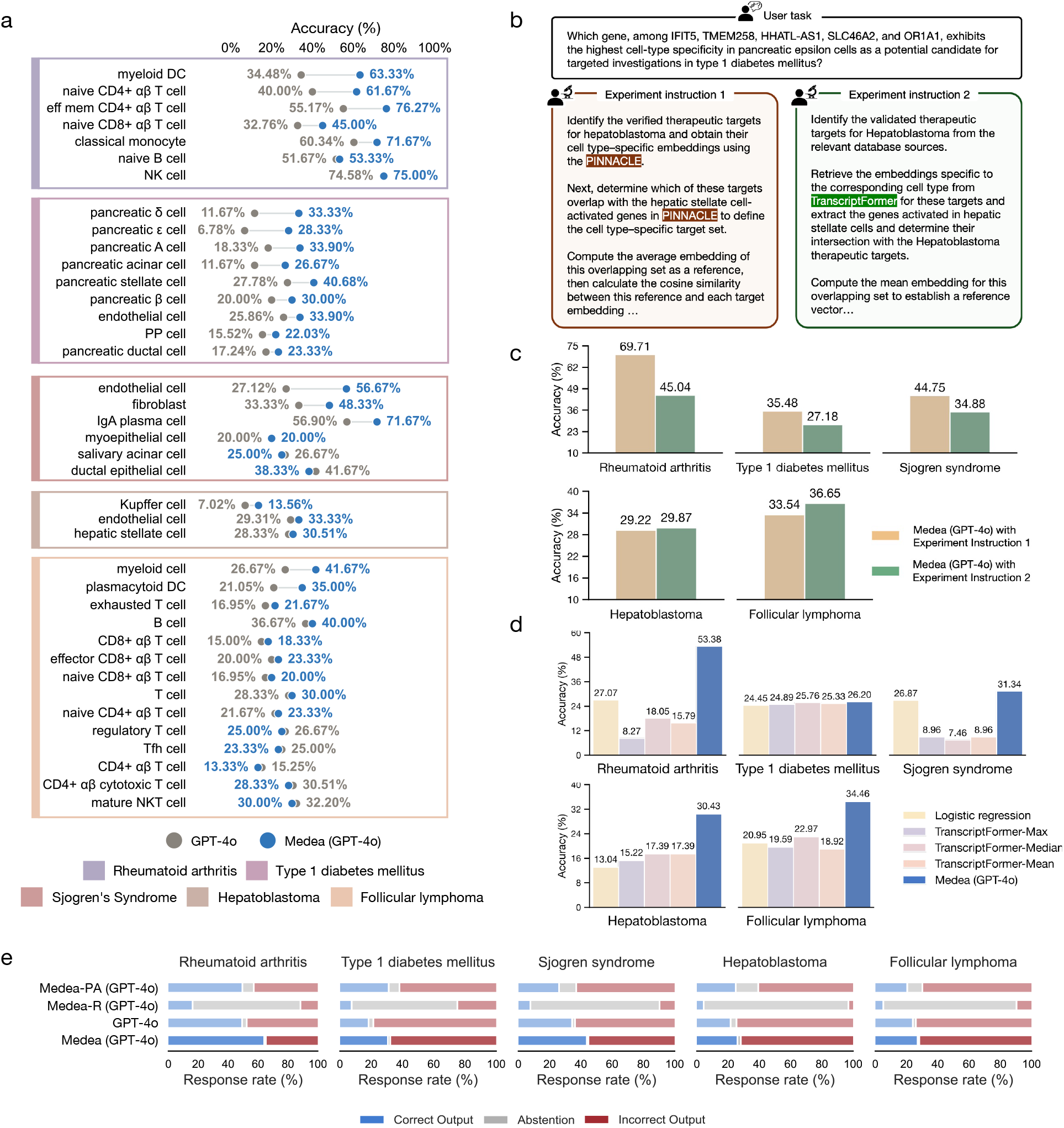
Medea performs cell type specific target nomination through multimodal agentic reasoning. **(a)** Accuracy of Medea (GPT-4o) and its backbone LLM (GPT-4o), stratified by disease and cell type, on the cell type specific target nomination benchmark. **(b-c)** Tool-constrained experiments in which Medea is instructed to use a specific single-cell foundation model (PINNACLE [26] or TranscriptFormer [27]) for linking the candidate genes to disease-relevant cell types, evaluating the impact of tool choice on performance. **(d)** Accuracy of Medea (GPT-4o) compared against TranscriptFormer as a standalone baseline across the five disease contexts. We evaluate TranscriptFormer in a zero-shot setting, ranking candidate targets by aggregated gene-embedding similarity (Max, Median, or Mean), and in a finetuned setting, where a logistic regression classifier is trained on the concatenated candidate embeddings (Methods 5.2). **(e)** Agent module ablations that quantify how planning, tool execution, and literature reasoning by Medea contribute to correctness and calibrated abstention. Medea-PA denotes the configuration that activates the ResearchPlanning and Analysis modules (Methods 1.2-1.3) to execute tool-based reasoning without literature synthesis. Medea-R denotes the configuration that activates only the LiteratureReasoning module (Methods 1.4). Medea denotes the full agent configuration that activates all four modules (i.e., ResearchPlanning, Analysis, LiteratureReasoning, MultiRoundDis-cussion).

### Verification and evidence reconciliation drive Medea’s performance

To quantify how Medea achieves cell type specific target nomination, we perform ablation analyses of its agentic modules. Because Medea can invoke different tools and agents, we isolate the contribution of each component by restricting Medea to specific subsets of tools and/or modules and measuring its accuracy on the omics objectives of nominating cell type specific therapeutic targets across five disease contexts (Figure 4b-e).

To assess how tool choice affects performance, we instruct Medea to use either PINNACLE [26] or TranscriptFormer [27] for the omics objectives of nominating cell type specific therapeutic targets (Figure 4b). Neither tool consistently outperforms the other when used by Medea (Figure 4c). With PINNACLE, Medea performs better for rheumatoid arthritis, type 1 diabetes mellitus, and Sjögren’s Syndrome. With TranscriptFormer, Medea performs comparably or better for hepatoblastoma and follicular lymphoma, respectively. These results show the strengths of different tools depending on the disease context, motivating agent access to a suite of complementary tools for completing omics objectives. Further, we find that Medea, equipped with Transcript-Former, generally outperforms TranscriptFormer in zero-shot and finetuned settings (Figure 4d; Methods 5.2). This suggests that Medea is conducting complex analyses using tools in addition to TranscriptFormer for reasoning about candidate targets across cell type and disease contexts.

Based on the omics-based objective, Medea can activate different combinations of modules to complete the analysis. To quantify the contribution of each module, we compare: pretrained knowledge from the backbone LLM only (GPT-4o), the ResearchPlanning and Analysis modules (Medea-PA), the LiteratureReasoning module only (Medea-R), and the full agent (Medea). Using all agentic modules generally yields the highest accuracy and the lowest abstention rates (Figure 4e). For example, because the literature on cell type specific therapeutic targets is limited, Medea-R abstains most frequently, with an average abstention rate of 79.1% across five diseases, whereas the backbone LLM abstains less (1.8% on average). However, the backbone LLM also produces the highest rate of incorrect nominations across disease contexts: 49.1% in rheumatoid arthritis, 80.0% in type 1 diabetes mellitus, 65.3% in Sjögren’s Syndrome, 76.1% in hepatoblastoma, and 75.5% in follicular lymphoma. These ablations show that Medea’s agentic modules are complementary and improve the overall agent’s performance.

### Inferring synthetic lethality in cellular contexts

We evaluate Medea for synthetic lethality prediction in cancer cell lines (Figure 5a) [83, 84]. Synthetic lethality occurs when perturbing two genes together reduces cellular viability substantially more than expected based on perturbing either gene alone, reflecting a synergistic interaction between the two gene functions. Synthetic lethality can be exploited to design therapies that selectively kill cancer cells while limiting toxicity to normal cells [83, 84]. Using a multilineage CRISPR screen that identified synthetic lethal interactions in seven cell lines [85], we construct an open-ended benchmark for synthetic lethality reasoning (Methods 3.1). The bench-mark includes 793 synthetic lethal gene pairs and 1,586 non-synthetic lethal pairs across seven cell lines: MCF7 (breast adenocarcinoma), MCF10A (fibrocystic breast disease), MDAMB231 (triple-negative breast cancer), CAL27 (tongue adenosquamous carcinoma), CAL33 (tongue squamous cell carcinoma), A549 (non-small cell lung cancer), and A427 (non-small cell lung cancer).

**Figure 5:**
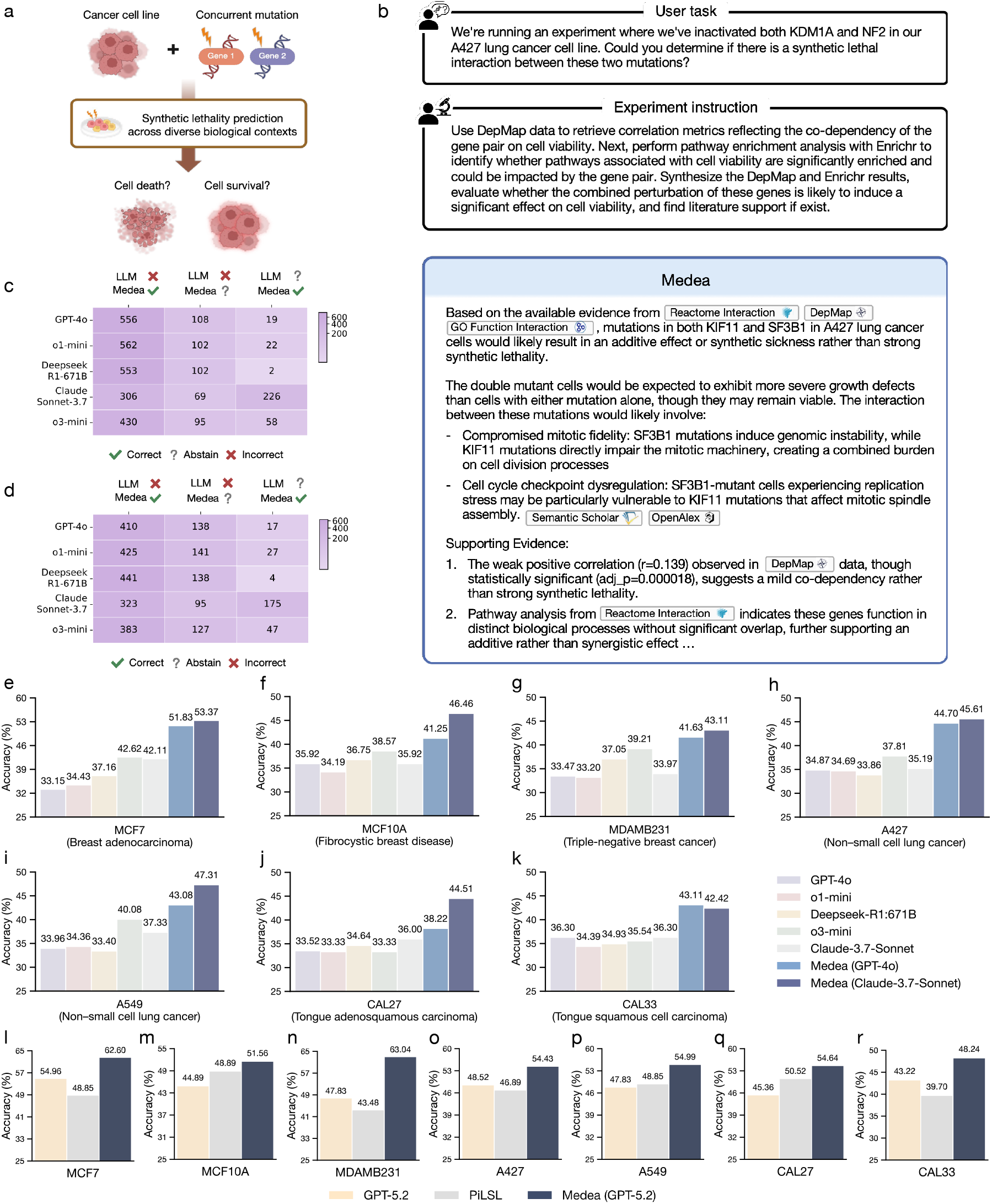
Agent reasoning about genetic interactions that affect cellular viability. **(a)** Given a pair of genes and a cell line context, Medea completes the omics objective of assessing whether the genes’ combined perturbation is consistent with reduced viability beyond single-gene effects (synthetic lethality; Methods 3). **(b)** Medea takes a natural language description of the omics objective (user instruction) and an optional experiment instruction, and produces an evidence-grounded response using tools (e.g., genetic dependency maps, pathway enrichment analyses) and literature evidence. **(c)** For each of five standalone LLMs, shown is the number of synthetic lethality analyses in which: (1) the LLM is incorrect and Medea is correct, (2) the LLM is incorrect and Medea abstains, and (3) the LLM abstains and Medea is correct. Medea uses a GPT-4o backbone. **(d)** Same as **(c),** with Medea using a Claude 3.7 Sonnet backbone. **(e-k)** Performance across seven cell lines of Medea and five LLMs on synthetic lethality reasoning. **(l-r)** Performance of Medea and PiLSL [111], a state-of-the-art machine learning model for synthetic lethality prediction, across seven cell lines.

Medea can activate all modules and tools, including tools that analyze DepMap gene co-dependency scores from CRISPR-Cas9 essentiality screens in cancer cells [86], biological pathways [29, 32, 87, 88], and molecular function datasets [33, 89, 90] (Methods 1.1). Across all seven cell line contexts, Medea yields higher accuracy than LLMs and reasoning models on inferring synthetic lethal interactions (Figure 5e-k). Medea achieves stronger accuracy by up to 20.2% in MCF7 (*p*-value = 0.0002 using McNemar’s test [60]; Figure 5e), 12.3% in MCF10A (*p*-value = 0.0012; Figure 5f), 9.9% in MDAMB231 (*p*-value = 0.0220; Figure 5g), 11.2% in CAL27 (*p*-value = 0.0145; Figure 5j), 8.7% in CAL33 (*p*-value = 0.0018; Figure 5k), 13.9% in A549 (*p*-value *<* 0.0001; Figure 5i), and 11.8% in A427 (*p*-value *<* 0.0001; Figure 5h). We observe performance gains compared to LLMs and reasoning models regardless of Medea’s LLM backbone. While Medea (GPT-4o) and Medea (Claude 3.7 Sonnet) have comparable performance in the MDAMB231 (Figure 5g), CAL33 (Figure 5k), and A427 (Figure 5h) cell line contexts, Medea (Claude 3.7 Sonnet) yields higher accuracy in MCF7 by 1.5% (Figure 5e), MCF10A by 5.2% (Figure 5f), CAL27 by 6.3% (Figure 5j), and A549 by 4.2% (Figure 5i).

Further, Medea, equipped with the latest LLM (GPT 5.2), outperforms PiLSL, a state-of-the-art SL prediction model, across all seven cell lines even if the LLM alone does not (Figure 5l-r; Methods 5.2). For example, in MDAMB231, Medea performs significantly better than its LLM backbone and PiLSL by 15.2% (*p*-value = 0.0002) and 19.6% (*p*-value = 0.0005), respectively (Figure 5n). Notably, Medea is able to achieve such strong performance even without leveraging PiLSL, as it is excluded from the tool space to assess Medea’s ability to predict synthetic lethality.

### Medea corrects errors and abstentions of LLMs

We test whether verification enables Medea to revise intermediate steps and convert unreliable LLM outputs into more reliable conclusions for omics objectives, which is important when prioritizing candidate perturbations for follow-up. We therefore focus on analyses where Medea is correct while an LLM used alone is incorrect or abstains (Figure 5c-d). Medea (GPT-4o) and Medea (Claude 3.7 Sonnet) correctly infer the viability outcome in at least 323 (13.6%) and 306 (12.8%) analyses, respectively, for which GPT-4o, o1-mini, Deepseek R1 671B, Claude 3.7 Sonnet, and o3-mini are incorrect (Figure 5c-d). In addition, for up to 175 (7.3%) and 226 (9.5%) gene pairs where an LLM abstains, Medea (GPT-4o) and Medea (Claude 3.7 Sonnet) correctly identify the interaction (Figure 5c-d). In these cases, Medea uses literature and data tools to re-evaluate the gene pair, check consistency with reported genetic screens, and prioritize interactions supported by convergent evidence, refining synthetic lethality inference beyond parametric knowledge alone.

Because false positives and false negatives can trigger unnecessary wet-lab experiments or missed therapeutic opportunities [84], we next analyze cases where an LLM is incorrect but Medea abstains. Medea (GPT-4o) and Medea (Claude 3.7 Sonnet) abstain in up to 141 (5.9%) and 108 (4.5%) analyses, respectively, when an LLM makes an incorrect prediction (Figure 5c-d). Although Claude 3.7 Sonnet has a relatively high abstention rate when used alone, it does not degrade the performance of Medea (Claude 3.7 Sonnet), indicating that Medea’s abstention behavior is not solely determined by the LLM backbone. Instead, abstention reflects the joint effect of the ResearchPlanning, Analysis, and LiteratureReasoning modules, which surface internal inconsistencies and choose not to commit to a synthetic lethal prediction. For example, Medea abstains when quantitative signals are weak or when pathway and empirical evidence do not corroborate the interaction (Supplementary Note 7). These results show that Medea can correct LLM errors and selectively abstain in uncertain cases of synthetic lethality inference.

### Predicting DNA damage-specific genetic interactions in yeast

Genetic interactions can be constitutive, detectable under standard growth conditions, or conditional, emerging only when cells are challenged by external stress [24]. Conditional interactions reveal functional buffering between pathways that is otherwise invisible at baseline, and are particularly informative for genome maintenance pathways whose activity is relevant mainly during DNA damage repair rather than during normal growth [91]. We therefore test whether Medea can reason about such context-dependent genetic interactions using two genotoxic agents that engage distinct repair pathways, bleomycin, which induces double-strand breaks repaired primarily through homologous recombination, and dimethyl sulfate, which induces base alkylation repaired primarily through base excision repair [25]; gene pairs that become synthetic lethal specifically under one agent therefore implicate redundant or buffering roles within the corresponding repair pathway rather than a general growth defect.

Given a yeast strain with concurrent mutations in a pair of genes, Medea is instructed to reason about the cell’s viability when exposed to either bleomycin or dimethyl sulfate (Figure 6a; Methods 3.3). Even without any experiment instruction, Medea is able to automatically select yeast-related tools to analyze the Alliance of Genome Resources [92], Saccharomyces Genome Database [93, 94], PomBase [95], STRING [96], and Costanzo SGA datasets [24, 91] (Figure 6a). We construct a synthetic lethality benchmark using a previously unpublished experimental screen that systematically maps genetic interactions in yeast. We perform high-density Epistatic MiniArray Profile (E-MAP) screening, which generates a 6,144-colony plate that profiles the genetic interactions between each query gene and genes in the full yeast single-deletion library (Methods 3.2). We experimentally screen 41 query genes, producing a 5, 806 × 41 matrix of gene pairs. Comparing the fitness of double-mutant strains against the expected fitness of the single mutants reveals the functional relationships between a pair of genes, with strong negative scores indicating synthetic lethality. By quantifying interaction scores with and without DNA-damaging treatment, we distinguish constitutive genetic interactions from those that emerge specifically under DNA damage stress. The benchmark includes 86 and 114 synthetic lethal gene pairs and 172 and 228 non-synthetic lethal pairs under two conditions, bleomycin (BLEO) and dimethyl sulfate (DMS), respectively.

**Figure 6:**
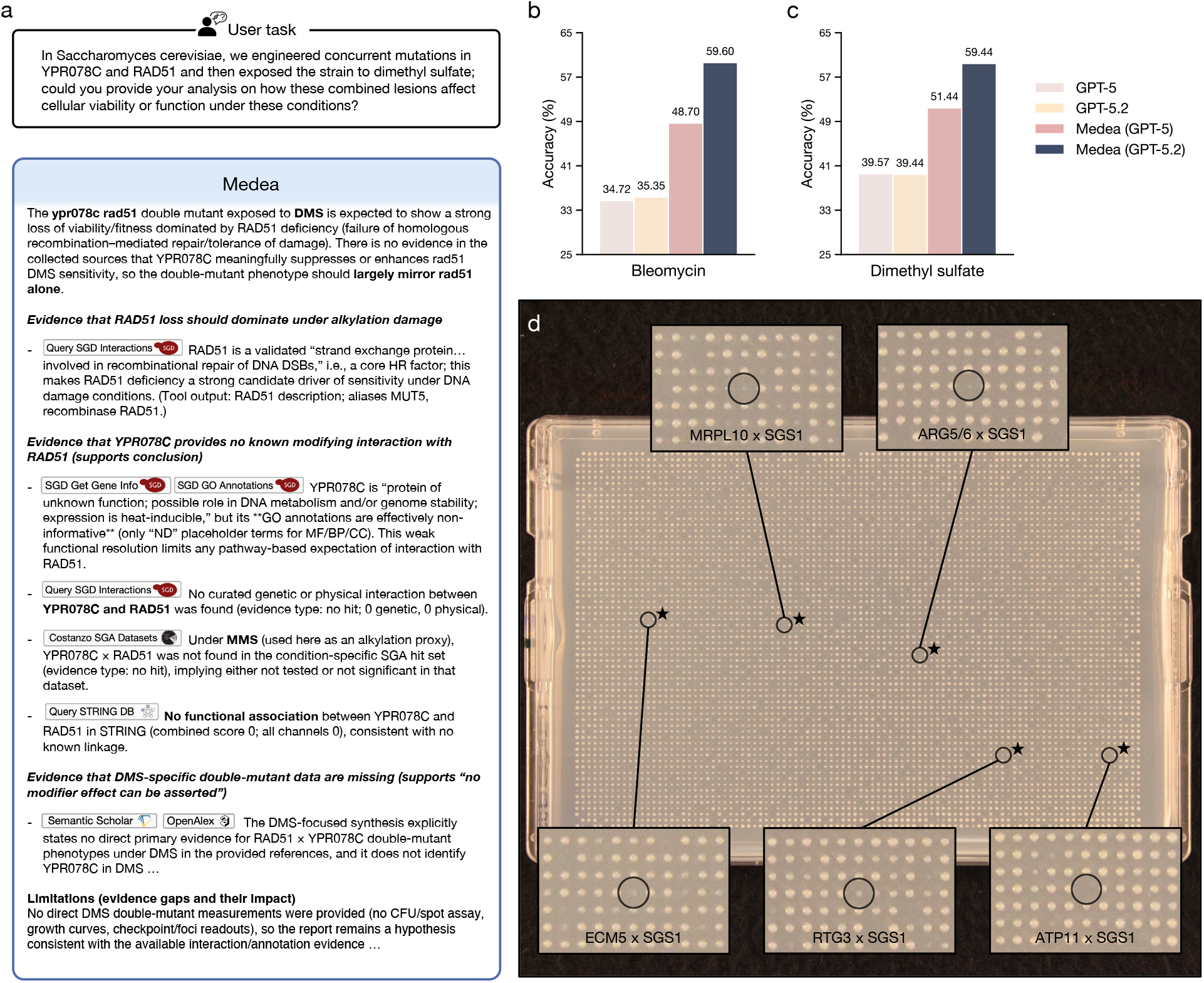
Experimental validation of synthetic lethal interactions in yeast. **(a)** Given a strain of *Saccharomyces cerevisiae* carrying concurrent mutations in a pair of genes, Medea reasons about the cell’s viability under exposure to bleomycin or dimethyl sulfate and returns an evidence-grounded report (Methods 3.3). Without an experiment instruction, Medea selects yeast-specific tools on its own, querying the Alliance of Genome Resources, SGD, PomBase, STRING, and the Costanzo SGA datasets. **(b-c)** Accuracy of Medea and its backbone LLMs (GPT-5, GPT-5.2) in predicting synthetic lethality in yeast under bleomycin (b; 86 synthetic lethal and 172 non-synthetic lethal pairs) and dimethyl sulfate (c; 114 and 228 pairs). Comparisons are head-to-head on analyses in which neither model abstains, so *n* differs across model pairs; accuracy over all non-abstention cases is reported in Supplementary Table 4. **(d)** A 6,144-colony plate from an Epistatic MiniArray Profile screen (Methods 3.2) profiling genetic interactions between SGS1 and a genome-wide library of yeast single-deletion strains, grown under dimethyl sulfate. Insets highlight five double knockouts predicted by Medea to be synthetic lethal with SGS1 under this condition: MRPL10, ARG5/6, ECM5, RTG3, and ATP11. Circles mark the array positions of these double mutants; the absence of a colony indicates loss of viability. A star denotes that the single knockout of each gene is viable (Supplementary Figure 4), so cell death reflects the combined perturbation rather than a single-gene fitness defect.

In a head-to-head comparison of non-abstention cases (i.e., neither model abstains), Medea outperforms its LLM backbone at predicting synthetic lethal interactions in yeast (Figure 6b-c). Under bleomycin, Medea yields higher accuracy than GPT-5 and GPT-5.2 by 14.0% (*n* = 193; *p*-value = 0.0001 using McNemar’s test [60]) and 24.3% (*n* = 99; *p*-value = 0.0015), respectively. Similarly, under dimethyl sulfate, Medea achieves stronger accuracy than GPT-5 and GPT-5.2 by 11.9% (*n* = 278; *p*-value = 0.0002) and 20.0% (*n* = 180; *p*-value = 0.0002). These performance gains are also observed when all non-abstention cases are considered (Supplementary Table 4).

We present a 6,144-colony plate from an E-MAP screen that profiles the genetic interactions between SGS1 and a genome-wide library of yeast single-deletion strains (Figure 6d). Visually and quantitatively, we validate Medea’s predictions of SGS1’s synthetic lethal interactions with MRPL10, ARG5/6, ECM5, RTG3, and ATP11. While single knockouts of these genes are viable (Supplementary Figure 4), our E-MAP screen shows that double knockouts lead to cell death.

### Treatment response prediction from tumor transcriptomes

We apply Medea to multimodal patient data, including clinical and transcriptomic profiles, to predict personalized response to immunotherapy (Figure 7a-b). Immunotherapy aims to treat cancer by activating the patient’s immune system against tumor cells; immune checkpoint inhibitors block proteins, such as CTLA-4 and PD-1, on T cells so that these cells can recognize and kill tumor cells [97]. However, biomarkers of treatment response are limited [98], and established ones, including tumor mutational burden (TMB) and features of the tumor microenvironment, are not consistently reliable predictors [6]. Using the IMvigor210 patient cohort [99], we construct an open-ended benchmark of 298 patients with bladder urothelial carcinoma treated with atezolizumab monotherapy (Methods 4). For each patient, Medea generates a report that includes a predicted responsiveness score and a rationale grounded in evidence relevant to the patient’s tumor transcriptomic profile, tumor microenvironment, and clinical profile (Figure 7a).

**Figure 7:**
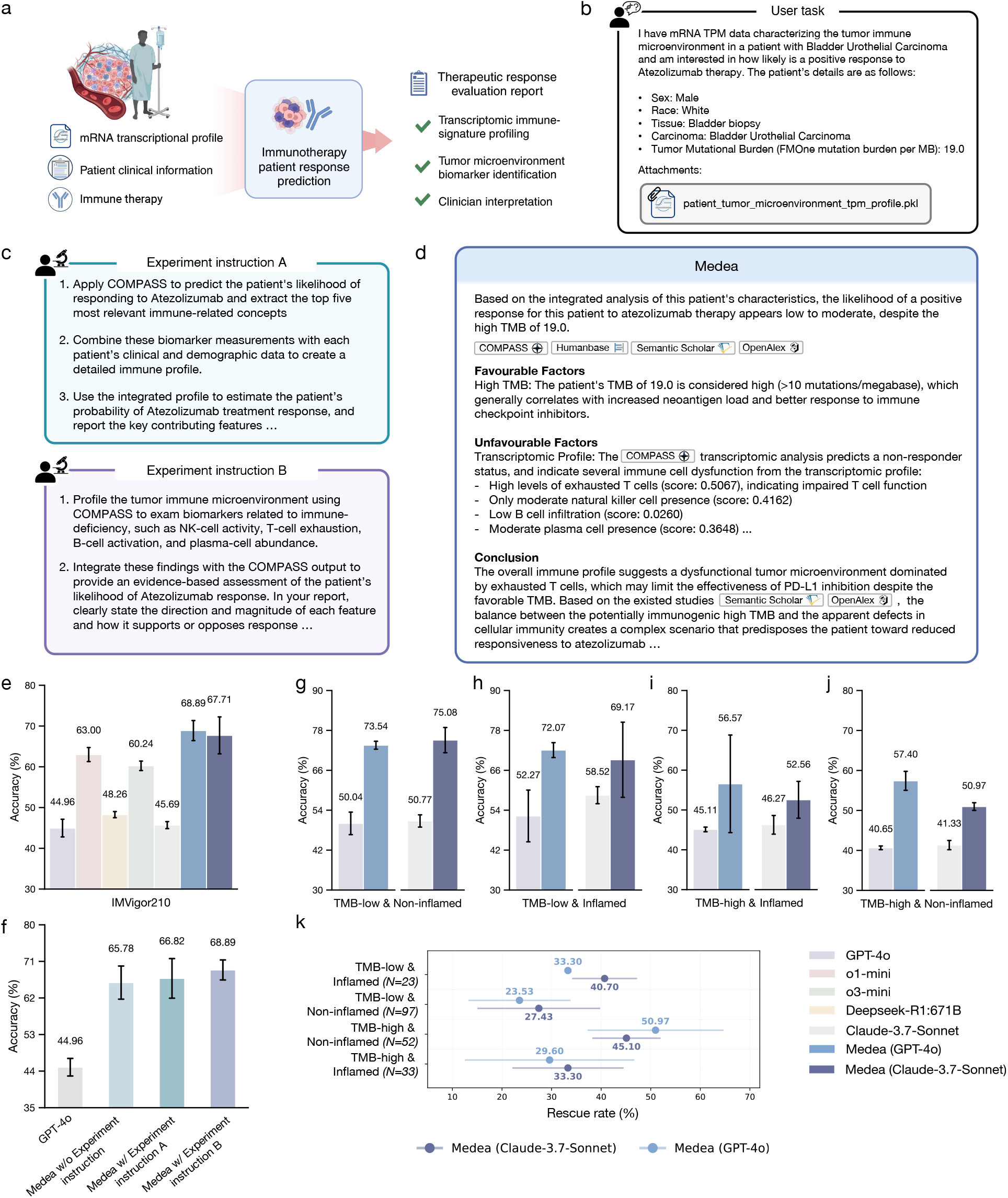
Multimodal reasoning for immunotherapy response prediction. **(a)** Given a patient’s clinical profile and tumor transcriptomic readout, Medea completes the omics objective of predicting immunotherapy response and returns a report with supporting evidence (Methods 4). **(b)** Multimodal inputs to Medea comprise clinical covariates and a tumor transcriptomic profile. **(c)** Medea accepts broad or detailed experiment instructions. **(d)** Medea grounds its report in evidence from an omics machine learning model (COMPASS [28]), tissue-specific protein networks [29], and literature [39, 40]. **(e)** Performance of Medea and five LLMs on predicting immunotherapy response. **(f)** Sensitivity of Medea (GPT-4o) and its backbone LLM (GPT-4o) to prompt variants: omics objective only, omics objective with a broad experiment instruction, and omics objective with a detailed experiment instruction. **(g-j)** Stratified performance of Medea (GPT-4o) and Medea (Claude 3.7 Sonnet) and their corresponding backbone LLMs (GPT-4o or Claude 3.7 Sonnet) on four clinical subgroups in the IMvigor210 cohort [99]. **(k)** Rescue analysis on IMvigor210 [99] patients for whom Medea correctly predicts the response label despite the incorrect prediction by the omics machine learning model [28]. Rescue rates are stratified by clinical subgroup.

Medea can activate all modules and tools, including those that query COMPASS, an interpretable machine learning model for immunotherapy response prediction [28]. Medea reasons over the biologically grounded signatures learned by COMPASS to explain its prediction, including immune biomarkers, and combines these signals with evidence from other tools to generate the final report (Figure 7a). We evaluate Medea under three experiment instruction settings: without explicit experiment instructions, with broad guidance (experiment instruction A), and with detailed step-by-step constraints (experiment instruction B) (Figure 7c). While experiment instructions A and B both direct Medea to use COMPASS for analyzing the patient’s transcriptomic profile, experiment instruction A asks Medea to interpret the top five immune-related concepts ranked by COMPASS whereas experiment instruction B specifies the concepts to interpret.

Under experiment instruction B, Medea outperforms LLMs in predicting treatment response (Figure 7d). Medea achieves up to 23.9% higher accuracy than LLMs (*p*-value *<* 0.0001 using McNemar’s test [60]). Consistent with cell type specific target assessment and synthetic lethality prediction (Figures 3-5), changing the underlying LLM used by Medea does not substantially affect performance on treatment response prediction (Figure 7e). In addition, varying or omitting the experiment instruction yields comparable performance, and Medea continues to outperform LLMs across these settings (Figure 7f). These results show that Medea is robust to differences in instruction specificity while completing the omics objectives of predicting treatment response from patients’ tumor transcriptomes.

### Interpreting treatment response from TMB and microenvironment

Immunotherapy response varies substantially across patients and tumor states [6, 100]. Tumor mutational burden (TMB) is often associated with improved response [6], but response also depends on features of the tumor microenvironment [101]. For instance, high TMB together with an inflamed microenvironment is more consistent with response than high TMB with a non-inflamed microenvironment [6]. Given this dependence on multiple interacting factors, we evaluate whether Medea can extract and reason over evidence about both TMB and the tumor microenvironment when inferring immunotherapy response.

We evaluate Medea (GPT-4o) and Medea (Claude 3.7 Sonnet) against their LLM backbones on treatment response prediction for patients across four subgroups: low TMB and non-inflamed (*N* = 97, with 16 responder and 81 non-responders; Figure 7g), low TMB and inflamed (*N* = 23, with 2 responders and 21 non-responders; Figure 7h), high TMB and inflamed (*N* = 33, with 15 responders and 18 non-responders; Figure 7i), and high TMB and non-inflamed (*N* = 52, with 21 responders and 31 non-responders; Figure 7j) tumor microenvironment. Medea achieves the highest accuracy across all groups. In contrast, standalone GPT-4o and Claude 3.7 Sonnet perform near random chance; for example, they tend to predict by default that patients with high TMB are responders. Further, Medea complements tool outputs by recovering false predictions. Across the four clinical subgroups, Medea correctly predicts response for patients that COMPASS mis-classifies (Figure 7k), with rescue rates of up to 51.0% in challenging settings such as patients with high TMB and non-inflamed microenvironments. These results indicate that integrating transcriptomic, clinical, and mutational profiles with literature evidence improves treatment response inference, and that Medea benefits from agentic modules that combine tumor microenvironment from transcriptomes (via analysis of a patient’s transcriptomic profile by the COMPASS tool [28]) with TMB and other biomarker features (via literature retrieval tools) (Supplementary Notes 1-5). To illustrate how Medea supports treatment response prediction, we present two patient vignettes from the IMvigor210 cohort [99] treated with atezolizumab. For each patient, Medea integrates clinical variables, tumor mutational burden, and tumor transcriptomes to infer treatment response. The first vignette is of a white male patient with bladder urothelial carcinoma with tumor mutational burden of 38.0 and a non-inflamed tumor microenvironment. Using the pre-treatment tumor biopsy transcriptome together with clinical and mutational features, Medea (GPT-4o) and Medea (Claude 3.7 Sonnet) predict response to atezolizumab. In this case, the Analysis and LiteratureReasoning pathways are concordant, concluding that the patient will respond based on COMPASS’s predicted responsiveness and immune signatures, including moderate-high cytotoxic T cell signal and interferon signaling, together with literature evidence linking treatment response to tumor microenvironment features and molecular subtype. GPT-4o and Claude 3.7 Sonnet used alone also predict response. The patient exhibits a partial response (RECIST [102]) after treatment. In a second vignette, we consider a white female patient diagnosed with bladder urothelial carcinoma, tumor mutational burden of 14.0, and an inflamed tumor microenvironment. Using the pre-treatment tumor biopsy transcriptome together with clinical and mutational features, Medea (GPT-4o) and Medea (Claude 3.7 Sonnet) predict non-response to atezolizumab. In this case, the Analysis and LiteratureReasoning modules provide conflicting evidence: Analysis predicts non-response based on COMPASS’s outputs [28], whereas LiteratureReasoning emphasizes the literature associating higher tumor mutational burden with better response to immune-checkpoint inhibitors [103]. After multi-round discussions by MultiRoundDiscussion, Medea predicts non-response by prioritizing the microenvironment signature and by noting that tumor mutational burden is not a consistently reliable biomarker when used alone, as factors like immune cell composition can dominate response. In contrast, GPT-4o and Claude 3.7 Sonnet used alone both predict that the patient will respond to treatment. The patient has progressive disease (RECIST [102]), consistent with Medea’s prediction of non-response.

## Discussion

A central challenge in therapeutic reasoning is that the same perturbation can produce different effects across diseases, cell types, tissues, and genetic backgrounds [1–3]. Medea addresses this problem through explicit context verification [71, 72]. In rheumatoid arthritis, for example, Medea distinguishes among closely related immune cell states, including naïve and effector-memory CD4^+^ *αβ* T cells, rather than collapsing analyses into broader lymphocyte categories. In follicular lymphoma, Medea prioritizes targets within disease-relevant B cell compartments and links these predictions to pathways associated with germinal center biology. These gains arise from preserving biological context throughout the agentic workflow, including the relationship between the biological question, the selected tools, and the datasets used for intermediate analyses.

Verification also improves performance during multi-step analyses [104, 105]. Agents using tools can fail when tool outputs are inconsistent or tool assumptions are violated [106, 107]. Medea introduces verification during both planning and execution, including checks for tool compatibility, parameter validity, and output consistency. Ablation analyses show that these components contribute substantially to performance. A literature-only configuration abstains frequently because literature evidence is often insufficient to complete a user task, whereas an LLM-only configuration produces substantially more incorrect conclusions despite low abstention rates. The full Medea configuration achieves the strongest overall performance.

Inferring synthetic lethality for over 200,000 gene-gene pairs in yeast extends our evaluation beyond human disease and cancer settings. Even without any predefined experiment instruction, Medea accurately predicts synthetic lethal interactions in yeast under bleomycin and dimethyl sulfate exposure using genetic interaction resources, pathway evidence, and literature retrieval adapted to the yeast context [24, 91]. By validating Medea’s predictions against epistatic miniarray profile screens, we additionally confirm that Medea can generate accurate predictions in settings where the experimental measurements are not yet available.

The immunotherapy analyses further illustrate the importance of integrating biological context during therapeutic reasoning. Medea combines tumor mutational burden with transcriptomic features of the tumor microenvironment, including interferon signaling, immune exhaustion, and antigen presentation programs [6, 101]. In several patient subgroups, Medea recovers correct predictions when standalone machine learning models or LLMs fail, suggesting that reconciliation across evidence sources can improve interpretation of heterogeneous clinical profiles. The relative robustness of Medea across prompt variants further indicates that verification and evidence reconciliation reduce sensitivity to differences in instruction phrasing [108].

Our study has a few limitations. First, the benchmarks rely on curated atlases, genetic dependency resources, and an immune checkpoint inhibitor naïve cohort, which may not capture the full diversity of tissues, perturbations, or clinical settings; additional cohorts will be needed to assess generalizability [28]. Second, some evaluations rely on LLM-based grading and human adjudication [109], which can introduce variability in assessment. Third, the analytical tools used by Medea encode assumptions about data preprocessing, cell type definitions, and pathway annotations. Although context verification reduces mismatches between tools and objectives, these assumptions can still influence downstream conclusions. Fourth, benchmark-based evaluations remain susceptible to information leakage because the training corpora of modern language models are incompletely characterized. The genome-wide yeast experiments address this limitation by validating predictions against experimental measurements that are not publicly available, but broader prospective validation will remain important. Finally, Medea’s consensus module depends on multiple language models and confidence-weighted reconciliation, which may still propagate correlated reasoning errors under some conditions [110].

Therapeutic hypotheses routinely move across diseases, model systems, and patient populations, but their validity depends on the biological context in which they are tested. Medea demonstrates that an agent combining context verification, tool-grounded execution, and evidence reconciliation can reason about targets within the cell states where they act, perturbations within a given genetic background, and treatment response within a patient’s transcriptomic and clinical context. These performance gains over LLMs, biomedical agents, and specialized machine learning models, together with experimental validation in a species and cellular context absent from Medea’s training and evaluation data, indicate that the capabilities transfer beyond the settings in which they were developed. As AI agents take on a larger role in assessing therapeutic hypotheses, verification and evidence reconciliation of the kind implemented in Medea will be necessary to keep agentic reasoning grounded in the biological context it is meant to represent.

## Acknowledgements

We thank NVIDIA for the helpful discussions on benchmarking DeepSeek models. M.M.L. is supported by the Berkowitz Family Living Laboratory at Harvard Medical School and the Clalit Research Institute. B.P.M., K.L., and T.I. are supported by NIH NCI U54 CA274502 and NIH NHGRI R01 HG012351. We gratefully acknowledge the support of NIH R01-HD108794, NSF CAREER 2339524, U.S. DoD FA8702-15-D-0001, ARPA-H Biomedical Data Fabric (BDF) Toolbox Program, Harvard Data Science Initiative, Amazon Faculty Research, Google Research Scholar Program, AstraZeneca Research, Roche Alliance with Distinguished Scientists (ROADS) Program, Sanofi iDEA-iTECH Award, GlaxoSmithKline Award, Boehringer Ingelheim Award, Merck Award, Optum AI Research Collaboration Award, Pfizer Research, Gates Foundation (INV-079038), Chan Zuckerberg Initiative, John and Virginia Kaneb Fellowship at Harvard Medical School, Biswas Computational Biology Initiative in partnership with the Milken Institute, Harvard Medical School Dean’s Innovation Fund for the Use of Artificial Intelligence, and the Kempner Institute for the Study of Natural and Artificial Intelligence at Harvard University. Any opinions, findings, conclusions or recommendations expressed in this material are those of the authors and do not necessarily reflect the views of the funders.

## Authors contributions

P.S., M.M.L, T.I., and M.Z. conceived the study. P.S. developed and implemented Medea. P.S., M.M.L., B.P.M., T.I., and M.Z. designed the experiments and interpreted the results. B.P.M., K.L., and T.I. conducted the genetic interaction screens in yeast and analyzed the results. P.S., M.M.L., and A.S. performed benchmarking and statistical analyses. M.M.L., A.S., and W.S. contributed agentic tools. M.M.L, V.G., and Y.H. retrieved, processed, and analyzed training and evaluation datasets. M.M.L. and S.G. contributed to the design of agentic modules, and Z.K. contributed to the agentic release and implementation. All authors discussed the research and contributed to the manuscript.

## Competing interests

T.I. is a co-founder, advisor, and holder of equity for Data4Cure and Serinus Biosciences, and he is an advisor and shareholder for Ideaya BioSciences and Eikon Therapeutics. The terms of these arrangements have been reviewed and approved by UC San Diego in accordance with its conflict of interest policies.

## Data availability

The model checkpoints, datasets, and agentic tools are available on Hug-gingFace at https://huggingface.co/datasets/mims-harvard/MedeaDB. The yeast EMAP screen is available at https://doi.org/10.6084/m9.figshare.32782446.

## Code availability

Project website is available at https://medea.openscientist.ai. The implementation of Medea and the code to reproduce the analyses in the paper are available on GitHub at https://github.com/mims-harvard/Medea.

## Online Methods

### 1 Overview of Medea

Medea is an omics AI agent for therapeutics discovery. Medea uses a tool space of databases, APIs, machine learning models, and agents (Section 1.1). It consists of a ResearchPlanning module (Section 1.2), an Analysis module (Section 1.3), a LiteratureReasoning module (Section 1.4), and a MultiRoundDiscussion module (Section 1.5).

### 1.1 Global tool space

Medea features a global tool space comprising 27 tools from 21 sources (17 databases/APIs and 4 machine learning models or agents). These tools are accessible to ResearchPlanning and Analysis modules throughout Medea’s runs.

**Retrieve Disease Targets** tool identifies and retrieves a disease’s protein targets through a two-step approach. Given a disease name, the tool first queries the EMBL-EBI API [30] to retrieve an Experimental Factor Ontology (EFO) ID for the disease. It then uses the EFO ID to interrogate the OpenTargets GraphQL API [31], extracting a list of associated protein targets based on specific evidence categories, such as genetic associations or approved therapeutic indications. By applying evidence-based filtering criteria, the tool outputs a curated list of protein targets.

**Load PINNACLE Embeddings** obtains pretrained cell type specific protein embeddings from PINNACLE [26]. PINNACLE integrates single-cell transcriptomics data with protein-protein interaction networks, resulting in 394,760 protein embeddings from 156 cell type contexts across 24 tissues. Given protein(s) and cell type context(s) of interest, this tool retrieves their corresponding cell type specific protein embeddings from PINNACLE. The final output is a dictionary containing context-specific protein embeddings, where keys are cell type contexts and values are cell type specific protein embeddings.

**Load TranscriptFormer Embeddings** retrieves cell type specific gene embeddings from TranscriptFormer [27], a single-cell transformer-based foundation model that models gene-gene interactions in single-cell transcriptomics data. The tool obtains precomputed Contextual Gene Embeddings (CGE) generated from single-cell atlases for rheumatoid arthritis [53], type 1 diabetes mellitus [112], Sjögren’s Syndrome [55], hepatoblastoma [56], and follicular lymphoma [57] (accessed via CELLxGENE [113]). Concretely, we perform TranscriptFormer inference on each cell type and disease combination using 16-bit mixed precision. As a result, we collect 2.1 million precomputed gene embeddings across 138 cell type and diseases. Given a list of gene names (official gene symbols or Ensembl IDs) along with disease and cell type contexts, the final output is a dictionary where keys are gene names and values are TranscriptFormer embeddings.

**Retrieve HumanBase PPI** retrieves tissue-specific protein-protein interaction networks, where nodes represent proteins and edges denote tissue-specific interactions annotated with confidence scores. Given a list of gene names and a tissue context, the tool first queries the Entrez API [114] to obtain the corresponding NCBI Entrez gene identifiers. It then calls the HumanBase RESTful API [29] to retrieve the relevant tissue-specific PPI network. From this network, an induced subgraph is generated based on the input gene list. The final output includes (i) a NetworkX [115] graph object representing the induced subgraph and (ii) a set of associations between the queried genes within the specified tissue context.

**HumanBase Co-Expression Analyzer** identifies genes that are co-regulated within a given tissue context. The tool accepts a gene list and a target tissue as inputs, and then queries the HumanBase API [29] to retrieve tissue specific co-expression interactions. Retrieved interactions undergo sequential quality control: raw correlations with coefficient ≤0.2 are excluded while those ≥0.7 are considered further. To ensure interpretability, the analysis does not rely solely on correlation strength: first, interactions lacking meaningful biological support are filtered out (evidence score ≤0.1); then, the top three most common evidence types behind the association are examined. The tool outputs a summary including co-expression network topology, network strength rating, ranked interactions with associated weights and supporting evidence, and a list of biological processes that are enriched in the co-expressed genes.

**HumanBase Protein Interaction Analyzer** identifies tissue-specific protein interactions by querying the HumanBase API, which integrates protein interaction data curated from BioGRID [116], IntAct [89], MINT [117], and MIPS [118]. Each protein pair is assigned a posterior probability weight based on experimental evidence. The tool retains edges with posterior probability ≥0.1 and prioritize those with ≥0.6 as high-confidence. Given a gene list, tissue name, and optionally the maximum number of interactions, the tool returns a summary of interaction types, confidence-weighted interaction pairs with supporting evidence, and a list of enriched biological functions.

**HumanBase Transcription Factor Analyzer** reconstructs tissue-specific regulatory networks for the genes of interest by analyzing transcription factor (TF)-gene relationships. Given a gene list, a target tissue, and optionally the maximum number of interactions, the tool queries the HumanBase API to retrieve a tissue-specific regulatory network derived from JASPAR motif predictions [29, 119]. TF-gene associations are identified through the scoring of binary motifs, with interactions filtered by evidence scores *>*0.1 to retain high-confidence predictions. The tool prioritizes TFs that may regulate the genes of interest in a given tissue context. The output includes the inferred TF-gene interactions, network connectivity scores, and enriched biological processes. **HumanBase microRNA Target Analyzer** evaluates how microRNAs modulate gene expression in specific tissue environments. Given a set of genes and a tissue context, the tool queries the HumanBase API for microRNA target interactions curated from MSigDB (c3:MIR) [90], and constructs the corresponding regulatory network. The tool aggregates evidence types across interactions, and summarizes primary evidence categories. It returns a summary of observed microRNA targeting patterns, key regulators, and associated functional pathways.

**HumanBase Perturbation Analyzer** extracts tissue-specific perturbation relationships from a given list of genes and tissue context. The tool retrieves perturbation data from MSigDB (c2: CGP) [90] via the HumanBase API, and retains only high-confidence associations (interaction weight ≥ 0.6) that are supported by evidence (evidence score *>* 0.1). Each association is evaluated for functional relevance by aggregating and interpreting the underlying evidence types. The tool returns: (1) perturbation response of gene pairs from MSigDB (e.g., gene A knockdown affects gene B expression), ranked by HumanBase interaction confidence scores; and (2) Gene Ontology biological process terms annotated to the query genes, retrieved from HumanBase’s curated annotations.

**Enrichr Gene Enrichment Analyzer** uses Enrichr RESTful APIs [87] to perform functional enrichment analysis for a given gene pair across multiple curated libraries: WikiPathways 2024 [88], Reactome 2024 [32], MSigDB Hallmark 2020 [33], and the 2023 releases of GO Molecular Function and GO Biological Process [34, 120]. Given a pair of genes, the tool retains terms with Benjamini–Hochberg adjusted *P* ≤ 0.05 as “enriched” terms (reported by Enrichr). It also records Enrichr’s combined score for each gene–term association, and constructs a bipartite graph where nodes are the two genes and the set of “enriched” terms. Each gene-to-term edge is weighted by Enrichr’s combined score, which integrates Fisher’s exact P with a z-score (*c* = ln(*P*) × *z*) [87, 121] to capture both statistical significance and effect size. The tool then aggregates the combined scores and counts shared terms to assign an overall confidence level, and labels the putative interaction mechanism (e.g., signalling, metabolic, regulatory, complex) based on the biological context of those terms. It returns a structured report containing: (i) a summary of the inferred gene–gene relationship in natural language, (ii) an overall confidence based on aggregated combined scores and shared-term counts, and (iii) the five most significant shared pathways or GO terms.

**WikiPathways Co-Pathway Inspector** examines whether two genes participate in shared biological pathways by querying the community-curated WikiPathways 2024 Human corpus [88]. Given a pair of genes, the tool uses the Enrichr REST API to retrieve all human pathways that contain each gene in the WikiPathways 2024 Human corpus, and intersects these sets to pinpoint pathways in which the genes co-occur. For every shared pathway, it derives an interaction-context label (e.g., signalling cascade, metabolic chain) from edge metadata, and a confidence score of the interaction based on node proximity, edge evidence, and annotation depth. The output is a structured list of pathway names, their interaction classifications, and their confidence scores.

**Reactome Co-Pathway Analyzer** queries the Enrichr REST API to identify genes involved in shared molecular reactions based on the Reactome Pathways 2024 database [32]. The tool determines functional associations between the given genes by analyzing whether two genes participate in the same reaction events (e.g., phosphorylation, complex formation, direct binding, enzymatic activity). It takes a pair of genes as input and returns a summary of overlapping biochemical interactions, including the classification of the molecular mechanism and a confidence estimate.

**Hallmark Co-Pathway Analyzer** identifies functional relationships between genes based on shared involvement in cancer hallmark processes, using the MSigDB Hallmark 2020 collection [33] obtained via the Enrichr REST API. Given two genes, the tool evaluates whether they co-occur in curated hallmark pathways associated with oncogenic processes (e.g., apoptosis, proliferation, metabolic reprogramming, immune signaling, DNA repair, angiogenesis). Outputs are a summary of shared hallmark processes, mechanism classifications, and confidence annotations.

**GO Molecular Function Checker** evaluates whether two genes share similar molecular functions by querying curated annotations from GO Molecular Function 2023 [34, 120]. Given a gene pair, the tool accesses the GO molecular function annotations via Enrichr REST API calls, and searches for convergent functional assignments, mechanism categories, and relative confidence scores. The outputs are a list of shared functional terms, associated weights, and relevant mechanism labels drawn from GO’s standardized molecular vocabulary.

**GO Biological Process Checker** analyzes the enrichment of biological processes among the input genes by querying curated annotations from GO Biological Process (GO-BP) 2023 [34, 87, 120]. Through Enrichr REST API calls, the tool tests each gene separately and intersects the significant GO-BP terms (Benjamini–Hochberg adjusted *p* ≤ 0.05). For every shared term, the tool records the GO ID and Enrichr’s combined score. It then summarizes the overlap into an overall confidence (e.g., number of shared terms, distribution of combined scores). The tool’s outputs are a list of shared GO-BP terms (names and GO IDs), per-gene statistics (adjusted *p*-value, combined score), mechanism labels, and an overall confidence summary.

**Retrieve DepMap Correlations** obtains pairwise co-dependency statistics using Chronos gene-effect profiles from the DepMap 24Q2 CRISPR dataset [35]. The tool accesses preprocessed Chronos gene scores derived from CRISPR-Cas9 knockout screens across 1,320 cancer cell lines. For each pair of genes, it calculates the Pearson correlation coefficients [122] and the BH-adjusted *p*-values between the Chronos scores of the two genes [123]. A Chronos dependency score of a gene estimates the likelihood that the gene is essential in a cell line (0 indicates not essential, and-1 suggests comparable essentiality to the median of all pan-essential genes) [124].

**Generate COMPASS Predictions** applies the COMPASS model to predict immunotherapy response based on a patient’s transcriptomic profile [28]. Given a patient’s transcriptomic profile (TPM) and cancer type, the tool calls COMPASS to (1) predict the scores of 44 biologically-grounded immune concepts, capturing immune cell states, tumor microenvironment interactions, and signaling pathways, and (2) predict the likelihood of the patient responding to immune checkpoint inhibitors (ICIs).

**Search Scientific Literature** retrieves and synthesizes relevant scientific literature to answer the user query. It queries the Semantic Scholar API to collect candidate papers [39], and filters for methodological soundness and direct relevance. The resulting set of papers is fed into an Open-Scholar reasoning module [41], which synthesizes their key findings into a concise response with citations to address the query. The output is either a structured dictionary containing the literature-grounded summary with inline citations or, if the search yields nothing suitable, an explicit note that no sufficiently relevant study is found.

**HPA Biological Processes Analysis** performs functional characterization of individual genes by querying the Human Protein Atlas (HPA) API [36–38], which integrates expression data from 44 human tissues, 10 cancer cell lines, 8 blood cell types, and 7 regions of the brain. The tool processes these data to generate three outputs. The tool retrieves tissue-specific expression data (nTPM), experimentally validated Gene Ontology (GO) annotations, and protein-protein interaction (PPI) datasets. It maps the retrieved GO annotations to ten canonical biological categories (e.g., cell cycle, apoptosis, metabolism) to stratify the gene’s involvement in core cellular mechanisms. Additionally, the PPI datasets retrieved from the HPA API are used to identify functionally similar proteins, quantifying similarity via a Jaccard index of shared biological process profiles. Finally, for genes with tissue-specific expression data, the tool calculates fold changes between cancer cell lines and healthy tissues to quantify the magnitude of differential expression. The final report includes the categorized GO annotations, functionally similar proteins identified by overlapping biological processes, and calculated fold changes across cancer cell lines for the queried genes.

**HPA Comparative Expression Analysis** performs comparative gene expression analysis on 10 cancer cell lines against healthy tissues [36–38]. For each gene, the tool retrieves normalized Transcripts Per Million (nTPM) values via the Human Protein Atlas (HPA) API, and performs several comparative analyses, including fold-change calculation, statistical significance assessment, and expression level categorization. The tool reports per-tissue nTPM, per-cell-line nTPM, and tissue-vs-cell-line fold-changes. By default, the tool summarizes results from a panel of 10 cell lines (HeLa, MCF-7, A549, HepG2, Jurkat, PC-3, RH-30, SiHa, U-251 MG, Ishikawa). Expression levels are binned into analysis-defined ranges (from very low *<*0.1 to very high ≥50 nTPM). To prioritize candidates, the tool calculates a differential expression score based on magnitude (*>*3-fold = high, *>*2-fold = moderate) and flags highly upregulated targets, returning a summary report containing HPA metadata and fold-change metrics.

***S. cerevisiae* Human Ortholog Mapper** identifies orthologous genes between *Saccharomyces cerevisiae* (yeast) and *Homo sapiens* (human) across three data sources using a priority-based merge. Given a yeast gene symbol or systematic ORF name (e.g., YER095W), the tool queries the Alliance of Genome Resources’s combined orthology dataset [92]. To address missing non-fungal homologs, the tool queries the Saccharomyces Genome Database (SGD) REST API [93, 94] and PomBase [95] for orthologs of *Schizosaccharomyces pombe*. The results from these three data sources are merged by matching on HGNC identifiers and gene symbols. Confidence levels are assigned based on the number of prediction methods that agree, according to the Alliance dataset (high: ≥8 methods, or ≥5 with best-score designation; medium: ≥3; low: otherwise). Orthologs found by both SGD/Alliance and PomBase are flagged as “cross-validated,” raising their confidence level. The tool accepts single-gene queries in either direction (yeast-to-human or human-to-yeast), and has a batch mode for processing gene lists from screens. Each output record includes the ortholog mapping, data sources, and associated confidence level.

**SGD Functional Complementation Retriever** queries the SGD REST API [93, 94] for experimentally-validated cases in which a yeast gene and a human gene can functionally rescue each other’s loss-of-function phenotype. Given a yeast gene symbol, the tool retrieves all of its complementation records curated in SGD. The output includes the direction of rescue (e.g., yeast complements human, human complements yeast), the strain background, experimental details, and the PubMed reference for each record.

**SGD Gene Annotation Retriever** retrieves gene annotations for a *Saccharomyces cerevisiae* gene from the SGD database [93, 94]. The tool queries the SGD features file via the gene symbol or systematic ORF name to extract the functional description, systematic name, gene type (e.g., ORF, tRNA, rRNA), and verification status (e.g., verified, uncharacterized, dubious). The tool falls back to the SGD REST API if the gene is absent from the file. The tool also returns aliases and the SGD identifier.

**SGD Curated Interaction Retriever** queries the SGD REST API for genetic and physical interactions sourced from BioGRID [116]. Given a pair of yeast genes, the tool retrieves all interactions for one gene and filters for records involving the other. For hub genes with a high interaction counts, the tool queries the low-degree genes first. Retrieved interactions are separated into “genetic” and “physical” categories based on the experiment type field, and genetic interactions are further classified as “aggravating” or “alleviating” based on the BioGRID annotation. The output includes classified interaction lists with the experiment type, annotation type (manually curated vs. high-throughput), phenotype, and PubMed reference for each record.

**STRING Functional Association Scorer** queries the STRING database [96] for integrated functional association evidence between two *Saccharomyces cerevisiae* genes. Given a pair of yeast gene symbols, the tool returns the combined confidence score and per-channel scores reported by STRING for seven evidence types: experimental, database, text-mining, co-expression, neighborhood, gene fusion, and co-occurrence.

**Costanzo Genetic Interaction Profiler** queries the Costanzo SGA datasets [24, 91] and collects the quantitative genetic interaction scores between two yeast genes. The tool accesses around 23 million gene pairs scored under standard growth conditions [24] and approximately 30,000 pairs profiled across 14 stress conditions, including DNA-damaging agents, protein-folding stressors, and signaling perturbations from [91]. For a given gene pair and condition, the tool returns the genetic interaction score (*ɛ*), p-value, and a classification as negative (aggravating), positive (alleviating), or neutral based on the sign and significance of *ɛ*.

**SGD Gene Ontology Annotation Retriever** collects the Gene Ontology (GO) annotations of a *Saccharomyces cerevisiae* gene across three GO domains (molecular function, biological process, and cellular component) via the SGD REST API [93, 94]. Given a *Saccharomyces cerevisiae* gene symbol, the tool retrieves and groups GO annotations by domain. Each annotation includes the GO term, GO identifier, evidence code, qualifier, and supporting reference. Annotations are deduplicated by term within each domain. The output is a structured dictionary organized by GO domain.

### 1.2 ResearchPlanning module

Given free-form user input, consisting of a user instruction and an optional experimental instruction, the ResearchPlanning module iteratively transforms it into *in silico* research plans (Figure 2b). It leverages three module-specific tools: ResearchPlanDraft, ContextVerification, and IntegrityVerification. *ResearchPlanDraft* generates a multi-step research plan that explicitly states the objectives of the analyses, the selected tools, and the tools’ expected inputs, parameters, and outputs; *ContextVerification* validates each biological entity and parameter choice against tool knowledge bases to confirm data availability and compatibility; and *IntegrityVerification* checks the specificity, technical feasibility, and logical consistency of the research plan.

**ResearchPlanDraft** transforms free-form user input into a structured multi-step research plan, which consists of sequential computational analyses that address the research objective. Each step specifies the analysis objective, selected tool, required inputs and parameters, and expected outputs. The procedure is: (1) apply an LLM to distill the objectives of the research plan based on the given instructions; (2) perform LLM-augmented retrieval (RAG) to identify relevant tools and obtain their metadata from the global tool space; and (3) assemble the objectives, relevant tools, and tool metadata into a coherent, stepwise research plan.

**ContextVerification** ensures that all biological entities and parameter choices in the plan are compatible with the available tools and datasets from the global tool space. The procedure is: (1) apply an LLM to extract the names and parameters of the tools specified in the plan; (2) verify the tools’ availability in the global tool space; (3) invoke each tool’s internal context checker function, when available, to validate the parameter choices in the plan; and (4) if the required context is missing or invalid, use an LLM to recommend a set of tool-supported alternatives.

**IntegrityVerification** acts as the final audit of the completeness, coherence, and logical soundness of the research plan. An LLM judge, guided by a rubric-style systems prompt, evaluates: (1) clarity of the plan; (2) tool-use fidelity, ensuring that each tool’s usage aligns with the documented functionality and parameter specification; (3) parameter precision, confirming that the inputs follow the documentation and best practices; (4) hallucination risk, flagging elements unsupported by the provided data or tool instructions; and (5) logical coherence, verifying that the steps are sequential, build upon each other appropriately, and follow a clear step-by-step progression toward addressing the objective. Failure on any criterion triggers a revision request (i.e., returning a message that asks for another refinement cycle to improve the plan). The tool outputs an evaluation report that summarizes the unmet criteria and provides concrete recommendations for revision.

### 1.3 Analysis module

The Analysis module translates the research plans from the ResearchPlanning module into well-documented, executable code for computational experiments. It coordinates four module-specific tools: CodeGenerator, AnalysisExecution, CodeDebugger, and AnalysisQualityChecker. *CodeGenerator* produces code snippets that implement the analyses specified in the research plan; *AnalysisExecution* runs the code in a sandboxed environment with resource and timeout controls, capturing standard output, error streams, and generated artifacts (e.g., tables, plots); *CodeDebugger* diagnoses runtime failures and revises code snippets using LLM-assisted debugging; and *AnalysisQualityChecker* performs post-execution evaluation, using structured criteria to assess correctness, reproducibility, parameter/provenance logging, and output informativeness against the user input and the research plan’s stated objectives. Altogether, the module operates through an autonomous iterative cycle, delivering executable code and *in silico* experimental results that address the user input and research plan without human intervention.

**CodeGenerator** uses a two-stage approach to generate executable code snippets for the analyses specified in the research plan. First, a ToolSelector component applies an LLM to identify the necessary tools and retrieve their associated metadata from the global tool space. Then, a separate LLM instance synthesizes code under a code-generation systems prompt that integrates the selected tools as outlined in the step-by-step research plan. Each generated code snippet undergoes a rubric-guided pre-execution check (e.g., syntax, interface compliance, parameter validity) using an LLM with a quality-check systems prompt; failures trigger bounded retries before returning a code snippet. This tool-aware code synthesis with iterative self-refinement is aligned with existing work on LLM tool-use and feedback-driven refinement [125, 126].

**AnalysisExecution** writes each code snippet to a temporary Python file and launches a Python subprocess with a 10-minute wait time to execute the code. Standard output and error streams are captured. Upon timeout, the tool captures the return code from the subprocess with the full traceback and logs attached for downstream analysis and debugging.

**CodeDebugger** is an LLM-based debugging tool. It analyzes the problematic code snippet and error messages, the user instructions, the research plan, and the tool specifications to identify the root cause. The CodeDebugger then generates corrected code snippets that address the errors while maintaining alignment with the research objectives and tool constraints, enabling iterative refinement until successful execution is achieved.

**AnalysisQualityChecker** evaluates the scientific validity and completeness of the successfully executed code using an LLM judge [109]. The tool analyzes the generated code and its execution outputs against the user instruction and research plan, examining code informativeness (i.e., outputs provide meaningful insights that are relevant to the research question), logical correctness (i.e., proper implementation of analytical workflows), and alignment with research objectives. The evaluation process uses structured LLM prompts that assess the quality of the analysis, and provides feedback in the tool’s output for iterative refinement if quality standards are not met.

### 1.4 LiteratureReasoning module

The LiteratureReasoning module performs autonomous retrieval, relevance appraisal, and synthesis of peer-reviewed publications to produce literature-grounded analysis using three module-specific tools: LiteratureSearch, RelevanceVerification, and OpenScholarReasoning. *Literature-Search* executes structured searches across academic journal indices (e.g., Semantic Scholar [39], OpenAlex [40]) to collect candidate papers. *RelevanceVerification* performs study quality screening, scoring each paper for topical relevance and study type and returning a compact evidence profile per paper. *OpenScholarReasoning* synthesizes the screened set of papers into a literature-grounded report to address the research objective in user instruction.

**LiteratureSearch** conducts literature searches across Semantic Scholar [39] and OpenAlex [40] databases. The process involves: (1) using an LLM to extract domain-specific search terms from the user instruction; (2) conducting literature searches in parallel with Semantic Scholar and OpenAlex tools; (3) aggregating the retrieved papers and their metadata, and performing deduplication based on exact title matching and DOI comparison; and (4) returning a curated collection of paper abstracts, citation counts, publication metadata, and source attribution for downstream processing.

**RelevanceVerification** systematically assesses the relevance of a paper to the research objective using an LLM. For each retrieved publication, the tool analyzes its title, abstract, and metadata to assess study characteristics (e.g., species, disease, cell type context), and generates a binary relevance label and a detailed relevance summary.

**OpenScholarReasoning** synthesizes evidence from the curated collection of papers using the OpenScholar framework [41] with integrated reranking capabilities. The process involves: (1) initializing a BGE reranker model [127] to identify and prioritize the most informative passages via semantic similarity to the user instruction; and (2) using OpenScholar [41] to generate a literature-grounded report (with proper citation formatting) that addresses the user instruction.

### 1.5 MultiRoundDiscussion module

The MultiRoundDiscussion module conducts a multi-round deliberation over the outputs of the Analysis module, LiteratureReasoning module, and backbone LLM to generate a consensus that addresses the user input. It adapts a ReConcile-style panel discussion [47] with standardized inputs, independent judgments, weighted reconciliation, iterative debate, and final synthesis.

Before deliberation, each module’s output is normalized into an evidence-grounded argument via a template with a fixed token budget and unified format. The deliberation has four main stages: (1) independent evaluation by each LLM panelist, (2) preliminary consensus among the panelists, (3) iterative discussion between the panelists until consensus is achieved, and (4) summarization of the panelists’ decision. In the first stage, three distinct LLMs independently evaluate each 300-word argument against the evaluation criteria (e.g., evidence strength, logical consistency, scientific rigor) to generate structured responses containing their preferred argument (i.e., from the Analysis module, LiteratureReasoning module, or backbone LLM), explanations, and confidence scores. The verdict of each LLM is saved in a JSON format. The three LLMs are configurable; by default, we use the backbone LLM, Gemini-flash-2.0 [128], and o3-mini [129]. In the second stage, preliminary consensus is determined by aggregating the verdicts from stage one via confidence-weighted voting using each panelist’s confidence score, which ranges from 0 to 1. If consensus is not achieved in stage two, the module initiates *R* (default is 2) rounds of debate. For each round, every panelist receives a debate prompt containing the arguments from each module and an audit trail summarizing the verdicts from the other panelists. Panelists reassess and update their own verdicts. When the panelists reach a consensus, the backbone LLM synthesizes the final response.

### 1.6 Implementation details

#### Base LLM configuration

Medea can be flexibly integrated with any LLM that supports function-calling and reasoning. Here, our experiments utilize Azure OpenAI GPT-4o (v2024-11-21; knowledge cutoff Sep 30, 2023) [130] and Claude Sonnet 3.7 (knowledge cutoff Feb 2025) [131] as backbone LLMs. Sampling temperatures are set to 0.4 for agent reasoning, 0.6 for tool invocations, and 1.0 for panel discussion modules; the remaining parameters are set to the default settings. For the yeast synthetic lethality experiments (Section 3.2-3.3) and the comparison against PiLSL (Section 5.2), we use Azure OpenAI GPT-5 (v2025-08-07; knowledge cutoff Sep 2024) and GPT-5.2 (v2025-12-11; knowledge cutoff Aug 2025) as backbone LLMs to evaluate Medea with more recent models.

#### AgentLite framework

Medea is developed using the AgentLite framework (v0.1.12) [132]. Each agentic module operates independently using the ReAct [133] architecture, performing an observe-act-reflect cycle for iterative tool-augmented reasoning. Individual modules maintain their own task-specific prompting schemas and memory modules for short-term trajectory tracking and long-term observation retrieval. As such, Medea’s architecture enables separation between reasoning logic, tool interaction, and task specialization.

### 2 Therapeutic target nomination

Nominating therapeutic targets requires reasoning about the candidate gene/protein in the disease and cellular contexts of interest. This is a biological question-answering task (Supplementary Table 1). We select five disease atlases from CELLxGENE [113, 134] for rheumatoid arthritis (RA) [53], type 1 diabetes mellitus (T1DM) [54], Sjögren’s syndrome (SS) [55], hepatoblastoma (HB) [56], and follicular lymphoma (FL) [57]. We process these disease atlases to identify disease-specific marker genes for each cell type (Section 2.1). With these marker genes and OpenTargets, we construct a benchmark dataset for therapeutic target nomination (Section 2.2).

### 2.1 Single-cell disease atlas processing

We process disease atlases from CELLxGENE [113, 134] using the standard scanpy [135] pipeline. First, we remove cells with fewer than 200 expressed genes. To control cell-level quality across the disease atlases, we apply thresholds on mitochondrial read fraction and detected gene count. For T1DM, we use a mitochondrial threshold of 25% and a gene-count cap of 6,000; for RA, 20% and 800; for SS, 0% and 3,000; for FL, 15% and 5,000; and for HB, 8% and 8,000. Next, we filter out genes that are expressed in fewer than 3 cells and that are among the 10% least variable genes. Then, we normalize the total UMI counts to a scaling factor of 10,000 reads per cell, apply log normalization, and scale each gene to unit variance (and clip values exceeding 10 standard deviations). We also filter out genes with missing or duplicated NCBI IDs, Entrez IDs, or gene symbols. To evaluate the processing quality, we perform UMAP dimensionality reduction on the processed data. We visualize cell clustering with respect to known biological and technical metadata, including cell type, donor identity, and other dataset-specific annotations. These embeddings allow us to assess whether biologically meaningful groupings are preserved while identifying potential batch effects, indicated by clustering due to technical rather than biological factors. Finally, to identify disease-specific marker genes for each cell type, we perform differential expression analysis using a one-vs-all Wilcoxon rank-sum test. For each cell type within a given disease context, expression is compared against all other cells belonging to different disease statuses. This approach identifies genes that are specifically enriched in particular cell type and disease combinations.

### 2.2 Dataset construction

Dataset construction follows a three-step procedure. First, we analyze the processed single-cell disease atlases from CELLxGENE [113] to identify disease-specific cell marker genes, which are significant differentially expressed genes (Wilcoxon rank-sum test with Bonferroni correction at *α* ≤ 0.05) within a cell type and disease context [136, 137]. Second, we collect 2,415 disease-associated genes/proteins from the Open Targets Platform [31]. We keep genes/proteins with either a genetic evidence score *>*0 [138] or ChEMBL evidence score *>*0 [59]. Third, we define ground-truth cell type specific disease targets as the genes/proteins satisfying the criteria from both steps 1 and 2. For each cell type context, we sample one target (i.e., positive gene/protein) and four negative genes/proteins that do not meet the criteria to form five-gene candidate lists. Prompts are paraphrased with o3-mini-0131 (temperature = 1.0) under a “biologist” role. Using three random seeds, we generate 20 samples per cell type per disease, producing 2,400 analyses: 420 for RA, 600 for T1DM, 360 for SS, 180 for HB, and 840 for FL (Figure 3b; Figure 4b; Supplementary Table 1).

### 3 Synthetic lethality prediction

Predicting synthetic lethality (SL) requires reasoning about genetic dependencies in a certain cellular context to infer whether perturbing two genes together reduces cellular viability substantially more than perturbing either gene alone [83, 84]. This is an open-ended reasoning task.

### 3.1 Human SL dataset construction

We curate experimentally-validated (via combinatorial genetic screening [85]) SL gene pairs and matched negative gene pairs. Gene pairs are collected from seven cell lines (six tumor-derived and one non-tumorigenic control from distinct genomic contexts) to capture diverse genomic dependencies [139]: *KRAS* gain-of-function mutations (MDAMB231, A427, A549), *PIK3CA* gain-of-function mutations (MCF7, CAL33), *TP53* mutations (MDAMB231, CAL27, CAL33), and no such mutations (MCF10A) [85]. These cell lines represent contexts in which cancer therapeutic targets have been extensively characterized using CRISPR screens [140]. For each positive gene pair in a specific cell line, (gene*_a_,* gene*_b_,* cell*_x_*), two negative gene pairs are generated via random substitution from a pool of 9,987 reported non-SL pairs, (gene*_a_,* gene*_c_,* cell*_x_*) and (gene*_b_,* gene*_d_,* cell*_x_*). The substituted genes (gene*_c_* and gene*_d_*) are experimentally-validated to not have a synthetic lethal interaction with the positive genes (gene*_a_* and gene*_b_*) in any of the cell line contexts [85]. This strategy preserves structural similarity to the positive pairs while ensuring non-lethality. With these triplets of gene pairs and cell line context, we construct an open-ended reasoning benchmark for context-specific SL prediction (Figure 5b; Supplementary Table 1). Prompts are paraphrased with o3-mini-0131 (temperature = 1.0) under a “biologist” role. The final dataset contains 2,379 analyses: 171 SL and 342 non-SL in A427; 180 SL and 360 non-SL in A549; 60 SL and 120 non-SL in CAL27; 120 SL and 240 non-SL in CAL33; 61 SL and 122 non-SL in MCF7; 117 SL and 234 non-SL in MCF10A; and 84 SL and 168 non-SL in MDAMB231.

### 3.2 Screening synthetic lethal interactions in yeast

Genetic interaction profiles between query genes of interest and a genome-wide library of yeast single-deletion strains were measured by high-density E-MAP (Epistatic MiniArray Profile) screening, following a previously described ultrahigh-density single-plate protocol [141]. Briefly, a *Saccharomyces cerevisiae* single-deletion library (BY4741-derived YKO library; Dharmacon) was distributed across four 1,536-density source plates and condensed onto a single 6,144-colony plate (Singer ROTOR), producing one fixed array position per array-side knockout. Each query strain, a *MATα* single knockout in the second gene of interest, was mated against the 6,144 array; diploids were selected and sporulated, and haploid *MAT*a double mutants carrying both deletions were recovered by standard SGA selection on synthetic dropout media. Final selection plates were grown at 30^◦^C under one of two independent DNA-damaging conditions: selection media supplemented with dimethyl sulfate (DMS), a DNA-alkylating agent, or selection media supplemented with bleomycin (BLEO). Across the EMAP screen, 41 query genes were profiled: BDF1, BDF2, BLM10, CDC4, CDC73, CHD1, CUL3, DCK1, DNL4, DST1, DUN1, ELA1, ELC1, ELG1, ERCC4, HO (WT), IDP2, IRA1, JHD2, LST7, MEU1, MSH6, PHO23, PLC1, RAD2, RAD23, RAD51, RAD54, RAD57, RAD9, RPD3, RSC1, RSC2, SAK1, SET2, SGS1, SIN3, SSM4, TEL1, TEP1, and TOS3. Each query was screened in six biological replicate plates, with the exception of ELA1 and RSC2 in the bleomycin condition (three and four replicate plates, respectively); SGS1 was profiled in two independent batches (twelve replicate plates in total). Plates were imaged at five timepoints spanning the colony growth window (3, 6, 9, 12, and 15 h for the EMAP3 batch; absolute imaging times for the remaining batches were not recorded) with a digital camera (Canon EOS Rebel T3i; 18–55 mm lens). Raw RGB images were segmented and colony pixel areas extracted with a custom MATLAB toolkit (version R2017b; https://github.com/brazilbean/Matlab-Colony-Analyzer-Toolkit). Colony-size trajectories across the five timepoints were used to estimate normalized colony growth rates, from which single-and double-mutant fitness values were derived. Interaction scores (*π*), quantifying the difference between observed and expected double-mutant fitness, were then computed as previously described [142]. The statistical significance of each interaction was assessed from its z-score and a false-discovery-rate (FDR) correction across all tested gene pairs.

### 3.3 Yeast SL dataset construction

Following [142], we define SL and non-SL gene pairs by applying stringent thresholds to the scored pairwise interactions under BLEO and DMS conditions. A gene pair is labeled as SL if all three conditions are satisfied: FDR *<* 0.05, *Z <* −3, and *π* + 3*σ_pair_ <* 0. The FDR and Z-score cutoffs follow the validation criteria of [142], which requires that *p <* 0.05 and |*Z*| *>* 3 to separate true interactions from background. The condition *π* + 3*σ_pair_ <* 0 ensures that the interaction score remains negative even at the upper bound of estimation uncertainty (*>* 99% coverage). Non-SL pairs are defined by FDR *>* 0.9 and |*Z*| *<* 1.0, restricting to gene pairs whose double-knockout phenotype is indistinguishable from background. Under these criteria, 93% of scored gene pairs are excluded. The BLEO condition yields 139 SL pairs out of 206,377 valid pairs, and the DMS condition yields 204 out of 208,745. This reflects an SL rate of less than 1%, which is consistent with genome-wide estimates from [24]. For each SL pair under a specific condition, two matched non-SL pairs are sampled using the same strategy as in Section 3.1. We keep SL pairs whose sampled negative pairs satisfy the non-SL criteria (i.e., FDR *>* 0.9 and |*Z*| *<* 1.0). Prompts are paraphrased using the settings described in Section 3.1. The final dataset contains 600 analyses: 86 SL and 172 non-SL pairs in BLEO, and 114 SL and 228 non-SL pairs in DMS.

## 4 Immunotherapy response prediction

Predicting personalized immunotherapy response requires reasoning about each patient’s clinical features, genomic profile (e.g., tumor mutational burden), and tumor microenvironment [6, 100, 101]. This is an open-ended reasoning task on multimodal inputs.

### 4.1 Dataset construction

We construct the dataset using the IMvigor210 cohort (*N* = 298) [99], a single-arm phase II study of anti-PD-L1 atezolizumab in patients with metastatic urothelial carcinoma. For each patient, we create a user instruction containing the patient’s clinical metadata: tumor mutational burden (TMB) [143], demographics (sex, race), tissue source, and treatment regimen (Figure 7b; Supplementary Table 1). We create two additional prompt templates that contain the user instruction and an experiment instruction to analyze the patient’s transcriptomic profile (Figure 7c). Prompts are paraphrased with o3-mini-0131 (temperature = 1.0) under a “clinician” role. Using three random seeds, we generate 2,682 analyses (3 seeds × 298 patients × 3 prompt templates).

## 5 Evaluation of model outputs

We provide details about model evaluation, performance metrics, and statistical analyses.

### 5.1 LLM judge

We use an LLM-as-a-Judge (or LLM judge) framework [109, 144] to assess model outputs. Notably, we incorporate mechanisms for selective prediction [145, 146] to handle three special cases:

1. *Abstain*, where the model explicitly admits insufficient evidence or inconclusive analyses [147];
2. *Failed*, where the model does not return any substantive analysis; and (3) *None*, where the model systematically evaluates and rejects all candidates based on context-specific criteria.

#### Multiple choice evaluation

For therapeutic target nomination (Section 2; Supplementary Figure 1) and immunotherapy response prediction (Section 4; Supplementary Figure 2), the LLM judge examines the model output to classify the prediction as one of the predefined categories. For therapeutic target nomination, the LLM judge either provides the target gene name or classifies the output into one of three categories (Supplementary Figure 1): *Abstain*, *None*, or *Failed*. For immunotherapy response prediction, the LLM judge classifies the output into one of four categories (Supplementary Figure 2): *R* (responder), *NR* (non-responder), *Abstain*, or *Failed*.

#### Open-ended reasoning

The LLM judge uses structured prompt templates that provide label definitions with examples [148]. This approach mirrors established methodologies for research claim verification, where the models must discern whether the evidence supports or refutes a hypothesis [149]. For synthetic lethality prediction (Section 3), the LLM judge evaluates the model’s open-ended reasoning trace to classify it as one of four categories (Supplementary Figure 3): (1) *Synthetic lethality*, (2) *Non-SL*, (3) *Abstain*, or (4) *Failed*.

### 5.2 Model evaluation

Medea is benchmarked against five large language models (LLMs), one biomedical agent, and two machine learning models.

#### Large language models (LLMs)

We evaluate Medea against five state-of-the-art LLMs: GPT-4o [130], o1-mini [150], o3-mini [129], DeepSeek-R1:671B [151], and Claude-3.7-Sonnet [131].

#### Biomedical AI agents

CellVoyager is designed to reproduce single-cell analyses from scientific papers [20]. We assess its ability to nominate therapeutic targets for rheumatoid arthritis [26].

#### Machine learning (ML) models

We compare Medea against two state-of-the-art ML models. We evaluate TranscriptFormer, a single-cell foundation model [27], in nominating therapeutic targets across cell type and disease contexts (Figure 4d). We use the precomputed CGEs described in Section 1.1. In the zero-shot setting: For each analysis (or prompt) in which all five candidate targets have cell type specific gene embeddings from TranscriptFormer, we compute pairwise cosine similarity scores between the embeddings of the candidate targets and non-overlapping cell type specific disease targets (Section 2.2); aggregate the scores for each candidate target (via `mean`, `median`, or `max`); and nominate the most likely target based on the highest aggregated score. In the finetuned setting: We concatenate the five candidate genes’ embeddings and apply a classifier to predict the index of the most likely target (i.e., *y* ∈ [0, 4]). The classifier (i.e., logistic regression) is trained on the samples from the most common cell type context. Performance is calculated on the remaining samples for which Medea generates a prediction (Figure 4d).

PiLSL [111] is a graph neural network for predicting synthetic lethal interactions. Following its original GitHub repository, we train 5 checkpoints of PiLSL under the C1 evaluation scheme using the recommended hyperparameter settings: 3-hop enclosing subgraphs, 3 GCN layers, 64-dimensional embeddings, batch size of 512, Adam optimizer, learning rate of 5 × 10^−3^, and L2 coefficient of 10^−4^. For inference on our human SL dataset, we apply a sigmoid function and compute the mean probability across the 5 checkpoints. A gene pair with a mean score ≥ 0.5 is classified as SL. Since PiLSL does not consider cell line context, the same prediction label is assigned to the gene pair in any cell line. We evaluate PiLSL on analyses (or prompts) in which both genes are present in the knowledge graph. To avoid data leakage, we exclude the gene pairs in our human SL evaluation benchmark that are seen during the training of PiLSL. Of the 2,379 benchmark analyses, 117 analyses are excluded.

### 5.3 Accuracy

We adopt the selective prediction evaluation protocol [152, 153]. We define accuracy as the proportion of analyses for which the model output contains a definitive and correct answer (Section 5.1). Formally, given *N_total_* analyses, the accuracy (or selective risk) is calculated as:

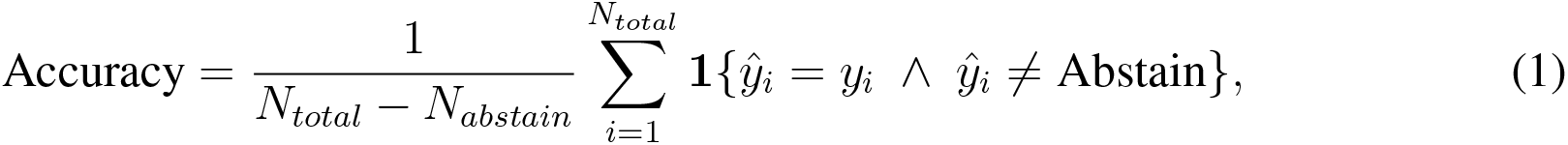

where **1**{·} is the indicator function that equals 1 when the predicted category matches the ground-truth label (case-insensitive) and 0 otherwise, and *N_abstain_* denotes the total number of analyses classified as *Abstain* (Section 5.1). *Abstain* cases occur when models explicitly acknowledge insufficient evidence or uncertainty in their reasoning. As abstentions are not incorrect predictions, we exclude them from accuracy calculations, decoupling correctness from coverage [145]. We additionally report the abstention rate *r_abstain_* = *N_abstain_/N_total_*.

### 5.4 Statistical significance testing

We use McNemar’s test [60] to evaluate the statistical significance between two models’ performance. We construct a 2×2 contingency matrix. Each entry *n_ij_* indicates the number of analyses in which Medea’s correctness is *i* ∈ {0, 1} and the other model’s correctness is *j* ∈ {0, 1}, where 0 represents correct and 1 represents incorrect. *n*_01_ denotes the number of analyses where Medea is correct but the other model is incorrect. *n*_10_ denotes the number of analyses where Medea is incorrect but the other model is correct. We exclude analyses where either model abstains (Section 5.1). The McNemar’s test statistic is calculated as:

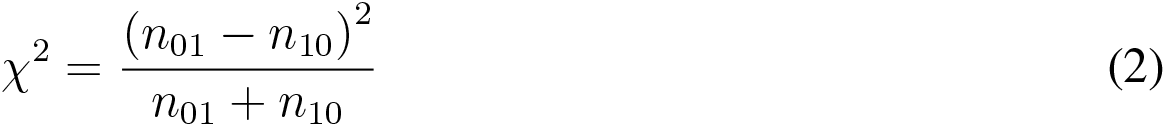

using the implementation from the statsmodels package [154].

## Notes

https://github.com/mims-harvard/MEDEA

https://doi.org/10.6084/m9.figshare.32782446

### Competing Interest Statement

The authors have declared no competing interest.

### Summary of Updates

Figure 4 revised and Supplymetory figure updated.

