## Supplementary Information for "Medea: An AI agent for therapeutic reasoning across biological contexts"

Medea is at https://medea.openscientist.ai
Medea code and benchmarks are at https://github.com/mims-harvard/Medea

#### Supplementary Notes

- **Supplementary Note 1**. Illustration of the omics objective on immunotherapy.
- **Supplementary Note 2**. Illustration of ResearchPlanning module outputs.
- **Supplementary Note 3**. Illustration of Analysis module outputs.
- **Supplementary Note 4**. Illustration of LiteratureReasoning module outputs.
- **Supplementary Note 5**. Final report generated by Medea.
- **Supplementary Note 6**. Evaluation of CellVoyager.
- **Supplementary Note 7**. Example abstention case from Medea.

#### Supplementary Tables

- **Supplementary Table 1**. Example user inputs from the three benchmark datasets for Therapeutic Target Nomination, Synthetic Lethality Prediction, and Immunotherapy Response Prediction.

- **Supplementary Table 2**. Accuracy (%) on the synthetic lethality prediction benchmark.
- **Supplementary Table 3**. Abstention rate (%) on the synthetic lethality prediction benchmark.
- **Supplementary Table 4**. Predicting synthetic lethality in *S. cerevisiae* under bleomycin and dimethyl sulfate conditions.

#### Supplementary Figures

- **Supplementary Figure 1**. Prompt template for instructing the LLM judge to evaluate the nominated therapeutic targets.
- **Supplementary Figure 2**. Prompt template for instructing the LLM judge to evaluate immunotherapy response predictions.
- **Supplementary Figure 3**. Prompt template for instructing the LLM judge to evaluate synthetic lethality predictions.
- **Supplementary Figure 4**. Single knockouts of SGS1 interaction partners are viable under dimethyl sulfate.

### Supplementary Notes

#### Note 1: Illustration of the omics objective on immunotherapy

The user input consists of user and experiment instructions. The example user instruction contains patient metadata and transcriptomic profile, and a question about the patient's likelihood of responding to Atezolizumab. The example experiment instruction directs the agent to analyze the patient's tumor microenvironment by applying COMPASS to the transcriptomic profile. Medea's outputs shown in Supplementary Notes 2-5 are generated using this example user input.

**User Instruction**

I have mRNA TPM data characterizing the tumor immune microenvironment in a patient with Bladder Urothelial Carcinoma. What is the estimated likelihood of a positive response to Atezolizumab therapy? The patient's details are as follows:

- Sex: Female
- Race: Caucasian
- Tissue: Kidney
- Carcinoma: Bladder Urothelial Carcinoma
- Tumor Mutational Burden (FMOne mutation burden per MB): 5.0

Attachment: patient-62fb13-ar-62fb138.pkl

**Experiment Instruction**

For patients with suspected immunodeficiency, begin by using the COMPASS tool. Start with an assessment of natural killer cell function, look for indicators of T cell exhaustion, and evaluate the activity of both B cells and plasma cells. Once you have calculated the COMPASS score, compare these results with transcriptomic markers of immune activation to identify those who may benefit from targeted therapies.

#### Note 2: Illustration of ResearchPlanning module outputs

The following research plan (generated via the user input from Supplementary Note 1) illustrates how the ResearchPlanning module decomposes the user and experiment instructions into a multi-step research plan. The ResearchPlanning module begins by summarizing the research objective from the user input and structuring the experiment as discrete steps. In this example, the ResearchPlanning module constructs a research plan to analyze a patient's tumor immune microenvironment using COMPASS [1]. The plan specifies evaluating NK cell activity, T cell exhaustion, B cell activation, and plasma cell abundance, with each step explicitly mapped to verified COMPASS concept names to ensure that the analysis uses functionally and semantically relevant parameters. The plan also integrates clinical metadata, including sex, ancestry, and tumor mutational burden. It concludes with a literature-grounded synthesis step, ensuring that the final response is supported by convergent statistical, mechanistic, and clinical evidence. This example demonstrates how the ResearchPlanning module transforms an analysis prescribed in natural language into a transparent, executable, and verifiable analysis plan.

Research plan generated by ResearchPlanning module

**Objective.** Determine the estimated likelihood of a positive response to Atezolizumab therapy for a Caucasian female patient with bladder urothelial carcinoma by analyzing tumor immune microenvironment transcriptomic data.

**Step 1: Load the patient's transcriptomic data**

**Action:**

- Load the patient's transcriptomic profile from the specified file path
- Verify the data format is compatible with the COMPASS tool

**API:** pandas
**API Method:** read_pickle

**Step 2: Analyze immune microenvironment using COMPASS**

**Action:**

- Process the patient's transcriptomic data through the COMPASS prediction tool
- Determine the predicted response to immune checkpoint inhibitor therapy
- Extract key immune-related concepts and their significance scores

**Tool:** compass_predict
**Description:** This tool leverages COMPASS, a research framework for concept feature extraction and Immune Checkpoint Inhibitor (ICI) response prediction, to analyze bulk transcriptomic data from a single patient.
**Import path:** from tool_space.compass import compass_predict
**Input Params:** df_tpm (pandas.core.frame.DataFrame) - A DataFrame containing the bulk transcriptomic data of a single patient (gene expression values in Transcripts Per Million format).
**Returns:**

- A boolean indicating the predicted responsiveness of the patient to Immune Checkpoint Inhibitor therapy (True for responder, False for non-responder).
- A ranked list of tuples, where each tuple contains an immune-related concept (as a string) and its corresponding significance score (as a float).

**Return Type:** bool, list[tuple[str, float]]

**Step 3: Analyze Natural Killer (NK) cell function**

**Action:**

- Extract NK cell-related scores from the COMPASS results using the standardized concept name 'NKcell'
- Evaluate the significance of NK cell activity in the tumor microenvironment
- Determine if NK cell function is normal, enhanced, or suppressed

**API:** pandas
**API Method:** DataFrame manipulation

**Step 4: Assess T cell exhaustion markers**

**Action:**

- Extract T cell exhaustion scores from the COMPASS results using the standardized concept name 'Exhausted_Tcell'
- Compare with other T cell metrics using standardized concept names ('CD8_Tcell', 'CD4_Tcell', 'Cytotoxic_Tcell')
- Determine the level of T cell exhaustion in the tumor microenvironment

**API:** pandas
**API Method:** DataFrame manipulation

**Step 5: Evaluate B cell and plasma cell activity**

**Action:**

- Extract B cell and plasma cell-related scores from the COMPASS results using standardized concept names ('Bcell_general', 'Plasma_cell')
- Assess the significance of B cell and plasma cell activity
- Determine if B cell and plasma cell functions are normal or altered

**API:** pandas
**API Method:** DataFrame manipulation

**Step 6: Analyze immune checkpoint interactions**

**Action:**

- Identify key immune checkpoint genes from the COMPASS results using the 'Immune_checkpoint' concept
- Focus on CD274 (official gene symbol for PD-L1, which is Atezolizumab's target) and related genes
- Analyze interactions between CD274 and other immune-related genes

**Tool:** analyze_comprehensive_interaction
**Description:** Performs comprehensive genetic interaction analysis by running all five Enrichr-based tools on a gene pair.
**Import path:** from tool_space.enrichr import analyze_comprehensive_interaction
**Input Params:**

- gene1 (str) - First gene symbol for comprehensive interaction analysis
- gene2 (str) - Second gene symbol for comprehensive interaction analysis

**Returns:** A dictionary with keys 'pathway', 'reactome', 'hallmark', 'function', and 'process'. Each value is a 4-element tuple: (summary, confidence, mechanisms, pathways).
**Return Type:** dict

**Step 7: Evaluate clinical significance of TMB**

**Action:**

- Analyze the significance of the patient's tumor mutational burden (TMB) of 5.0
- Correlate TMB with immune profile from COMPASS analysis
- Assess how TMB may influence response to Atezolizumab

**API:** pandas
**API Method:** DataFrame manipulation

**Step 8: Synthesize findings with scientific literature**

**Action:**

- Integrate all analysis results to determine likelihood of response to Atezolizumab
- Consider the patient's demographic information (Caucasian female)
- Evaluate how the immune profile and TMB together predict response to Atezolizumab
- Provide final assessment of response likelihood with supporting evidence

**Tool:** scientific_reasoning_agent
**Description:** Advanced literature synthesis tool powered by OpenScholar AI that queries 8M+ peer-reviewed research papers to provide evidence-based scientific reasoning.
**Import path:** from tool_space.agent_tool import scientific_reasoning_agent
**Input Params:** user_instruction (str) - Specific, focused scientific question or analysis request.
**Returns:** Comprehensive scientific analysis synthesizing relevant literature with evidence citations, mechanistic insights, clinical implications, and research recommendations.
**Return Type:** str

#### Note 3: Illustration of Analysis module outputs

The example experiment results reported by the Analysis module illustrate how it addresses the research plan laid out by the ResearchPlanning module (Supplementary Note 2). Based on the research plan, the Analysis module synthesizes executable code to perform the prescribed *in silico* analyses. The code is executed in a sandboxed environment with full provenance and error capture, producing reproducible computational results aligned with the plan. In this example, the Analysis module processes the patient's transcriptomic profile and applies COMPASS to predict a response outcome for Atezolizumab, accompanied by scores for immune-related concepts. The Analysis module additionally reasons about the interaction between CD274 (PD-L1) and CD8A (a surface marker on cytotoxic T cells) using Enrichr tools.

Experiment result reported by Analysis module

**Loading patient transcriptomic data from:** IMVigor210-62fb13-ar-62fb138.pkl
Successfully loaded data with 15673 gene expression values

**Analyzing immune microenvironment using COMPASS...**

Predicted response to Atezolizumab: RESPONDER

**Top 5 immune-related concepts:**

- Memory_Bcell: 1.3229
- Naive_Tcell: 1.1009
- Apoptosis_pathway: 1.0807
- Memory_Tcell: 1.0278
- Genome_integrity: 0.8705

**NK cell function assessment:**

- NK cell score: 0.3247 (ranked #37 of 42)
- Status: Normal

**T cell exhaustion assessment:**

- CD8_Tcell: 0.6995
- Exhausted_Tcell: 0.6603
- CD4_Tcell: 0.4771
- Cytotoxic_Tcell: 0.4091
- Exhaustion/Cytotoxic ratio: 1.61
- Status: High T cell exhaustion (unfavorable)

**B cell and plasma cell assessment:**

- Plasma_cell: 0.3451
- Bcell_general: 0.0291
- Status: Low B cell activity

**Analyzing immune checkpoint interactions...**

- Immune checkpoint score: 0.7513
- Analyzing CD274 (PD-L1) and CD8 T cell interactions...

**COMPREHENSIVE INTERACTION ANALYSIS**
Analyzing gene pair: (CD274, CD8A)

**PATHWAY ANALYSIS**

[PATHWAY] INSIGHT: Analyzing pathway interactions for CD274 + CD8A

[PATHWAY] INSIGHT: Strong evidence found: Cancer Immunotherapy By PD 1 Blockade WP4585 ($p=1.26 \times 10^{-6}$)

[PATHWAY] INSIGHT: Interaction detected: 1 shared terms, score=5.90

[PATHWAY] INSIGHT: Biological mechanisms identified: general pathway interaction

[PATHWAY] INSIGHT: Confidence 'low' based on: single shared term

[PATHWAY] INSIGHT: Pathway analysis complete: low confidence, 1 mechanism type

- Summary: Genes interact through 1 shared pathway(s) with low confidence
- Confidence: low
- Mechanisms: ['general_pathway']
- Key Evidence: ['Cancer Immunotherapy By PD 1 Blockade WP4585']

**REACTOME ANALYSIS**

[REACTOME] INSIGHT: Analyzing biochemical interactions for CD274 + CD8A

[REACTOME] INSIGHT: Interaction detected: 1 shared terms, score=2.74

[REACTOME] INSIGHT: Molecular mechanisms identified: general biochemical interaction

[REACTOME] INSIGHT: Confidence 'very_low' based on: single shared term

[REACTOME] INSIGHT: Reactome analysis complete: very_low confidence, molecular interaction detected

- Summary: Genes interact through 1 shared biochemical reaction(s) with very_low confidence
- Confidence: very_low
- Mechanisms: ['biochemical_interaction']
- Key Evidence: ['Adaptive Immune System']

**HALLMARK ANALYSIS**

[HALLMARK] INSIGHT: Analyzing cancer hallmark processes for CD274 + CD8A

[HALLMARK] INSIGHT: No shared terms detected between gene pair

[HALLMARK] INSIGHT: Genes do not co-regulate cancer hallmark processes

- Summary: Genes do not share cancer hallmark processes
- Confidence: none
- Key Evidence: None

**FUNCTION ANALYSIS**

[GO_FUNCTION] INSIGHT: Analyzing molecular functions for CD274 + CD8A

[GO_FUNCTION] INSIGHT: No shared terms detected between gene pair

[GO_FUNCTION] INSIGHT: Genes have distinct molecular function profiles

**PROCESS ANALYSIS**

[GO_PROCESS] INSIGHT: Analyzing biological processes for CD274 + CD8A

[GO_PROCESS] INSIGHT: No shared terms detected between gene pair

[GO_PROCESS] INSIGHT: Genes participate in distinct biological processes

- Summary: Genes do not share biological processes
- Confidence: None
- Key Evidence: None
- Pathway interaction: low confidence
- Summary: Genes interact through 1 shared pathway(s) with low confidence

**Tumor Mutational Burden (TMB) assessment:**

- Patient TMB: 5.0
- TMB status: Low (generally unfavorable for immunotherapy)

**Synthesizing findings with scientific literature...**

**FINAL ASSESSMENT**

Patient characteristics: Caucasian female with bladder urothelial carcinoma

**Key findings from literature analysis:**

Atezolizumab has demonstrated clinical activity in urothelial carcinoma largely through its modulation of the PD-1/PD-L1 axis, which relies on a robust and primed immune environment to achieve tumor regression. Several biomarkers have been explored to predict response to immune checkpoint inhibitors...

**Estimated likelihood of positive response to Atezolizumab:** High ($> 60\%$)

#### Note 4: Illustration of LiteratureReasoning module outputs

The following example illustrates the literature search analysis by the LiteratureReasoning module to address the user instruction (Supplementary Note 1). Here, the report generated by the LiteratureReasoning module integrates evidence from the IMvigor130 trial on biomarker dependencies for atezolizumab response [2], FGFR3-driven immunosuppression [3], and macrophage–T-cell interactions influencing ICB efficacy [4]. These studies reveal missing biomarkers (e.g., PD-L1 expression, FGFR3 mutation status) for decision-making, leading the agent to conclude that the patient's likelihood of response is below average without additional profiling.

LiteratureReasoning `module output`

**Estimating Response Likelihood to Atezolizumab in Bladder Urothelial Carcinoma**

**Tumor Mutational Burden as a Predictor**

For the Caucasian female patient with bladder urothelial carcinoma and a tumor mutational burden (TMB) of 5.0 mutations/Mb, the likelihood of response to atezolizumab (anti-PD-L1 therapy) appears to be relatively low based on available evidence. The IMvigor130 trial data suggests that higher TMB ($>$ 10 mutations/Mb) is associated with better outcomes for atezolizumab monotherapy compared to chemotherapy [1]. With a TMB of 5.0, which falls below this threshold, the patient would not be in the group expected to have particularly favorable outcomes with atezolizumab monotherapy.

**PD-L1 Expression and Its Importance**

A critical factor that would significantly influence the response prediction is the patient's PD-L1 expression status, which was not mentioned in the case description. Data from IMvigor130 shows that PD-L1 IC2/3 (high expression) was associated with significantly longer overall survival for atezolizumab monotherapy versus chemotherapy [1]. Furthermore, the combination of PD-L1 IC2/3 and high TMB ($>$ 10 mutations/Mb) identified a patient subset (approximately 14% of biomarker-evaluable patients) with particularly favorable outcomes with atezolizumab monotherapy [1]. Without knowing the patient's PD-L1 expression level, a complete assessment is challenging.

**FGFR3 Mutation Status Considerations**

Fibroblast growth factor receptor 3 (FGFR3) mutation drives T-cell-depleted microenvironment in UC, which led to the hypothesis that FGFR3 mutation might attenuate response to ICB in patients with metastatic UC [2]. The PSM analysis indicated FGFR3-mutated patients had worse overall survival (OS) in comparison to FGFR3-wildtype patients (HR=2.11, 95% CI=(1.16 to 3.85), p=0.015) receiving atezolizumab. The median OS was 9.2 months (FGFR3-mutated) versus 21.0 months (FGFR3-wildtype). FGFR3-mutated patients had lower disease control rate than FGFR3-wildtype patients (41.0% vs 66.7%, p=0.023) [2]. The meta-analysis involving 938 patients with metastatic UC confirmed FGFR3 mutation was associated with worse OS after ICB (HR=1.28, 95% CI=(1.04 to 1.59), p=0.02) [2]. Single-cell RNA transcriptome analysis identified that FGFR3-mutated UC carried a stronger immunosuppressive microenvironment compared with FGFR3-wildtype UC [2]. FGFR3-mutated UC exhibited less immune infiltration and lower T-cell cytotoxicity [2]. Higher TREM2+ macrophage abundance in FGFR3-mutated UC can undermine and suppress the T cells, potentially contributing to the formation of an immunosuppressive microenvironment. Lower inflammatory-cancer-associated fibroblasts in FGFR3-mutated UC recruited fewer chemokines in antitumor immunity but expressed growth factors to promote FGFR3-mutated malignant cell development. FGFR3-mutated UC carried an abundance of malignant cells characterized by high hypoxia/metabolism and low interferon response phenotype. Inhibition of FGFR3 might activate the immune microenvironment, and the combination of FGFR inhibitor targeted therapy and ICB might be a promising therapeutic regimen in metastatic UC, providing important implications for UC clinical management [2].

Another important factor to consider is the FGFR3 mutation status, which was not specified. Research has shown that FGFR3 mutation can attenuate the response to immune checkpoint blockade in metastatic urothelial carcinoma by driving an immunosuppressive microenvironment. The propensity score matched analysis from the IMvigor210 trial indicated that FGFR3-mutated patients had worse overall survival compared to FGFR3-wildtype patients receiving atezolizumab (median OS of 9.2 months versus 21.0 months) [2]. FGFR3-mutated patients also had a lower disease control rate than FGFR3-wildtype patients (41.0% versus 66.7%) [2].

**Immune Microenvironment Factors**

The tumor immune microenvironment is another critical determinant of response. Single-cell RNA sequencing studies have identified that immunosuppressive macrophages and exhausted T-cell subpopulations are enriched in the basal subtype of urothelial carcinoma and show enhanced interactions that may impact immunotherapy response [3]. The presence of a high fibroblast TGF-$\beta$–response signature (F-TBRS) has been associated with inferior overall survival with atezolizumab monotherapy [1], suggesting that stromal characteristics can influence treatment outcomes.

**Molecular Subtyping Considerations**

FGFR3 mutation drives T-cell-depleted microenvironment in UC, which can attenuate response to immune checkpoint blockade (ICB) in patients with metastatic UC [2]. Compared to FGFR3-wildtype patients, those with FGFR3-mutated metastatic UC have worse overall survival and lower disease control rates when receiving atezolizumab. Single-cell RNA transcriptome analysis has identified that FGFR3-mutated UC carries a stronger immunosuppressive microenvironment with less immune infiltration and lower T-cell cytotoxicity. Additionally, PD-L1 expression, tumor mutational burden (TMB), and T-effector gene expression have been associated with clinical outcomes in patients treated with atezolizumab [1]. Specifically, a combination of PD-L1 IC2/3 and high TMB identified a patient subset with particularly favorable outcomes when treated with atezolizumab monotherapy [1]. Both immunosuppressive macrophages and exhausted T-cell subpopulations have been found to be enriched in the basal subtype of urothelial carcinoma [3].

Molecular subtyping of the tumor would provide additional insights into the potential response. Studies have shown that the tumor's molecular subtype (basal versus luminal) can influence the immune microenvironment and, consequently, the response to immunotherapy [3]. The development of a gene expression signature (Macro-C3 score) capturing immunosuppressive macrophages has been shown to predict outcomes better than the currently established subtypes [3].

**Conclusion**

Given the patient's TMB of 5.0 mutations/Mb (below the favorable threshold of $> 10$) [1], without knowing PD-L1 expression status, FGFR3 mutation status [2], or molecular subtype [3], the estimated likelihood of a positive response to atezolizumab appears to be below average. For a more accurate prediction, additional biomarker testing would be recommended, including PD-L1 expression [1], FGFR3 mutation analysis [2], and comprehensive molecular profiling [3]. This approach would allow for a more personalized assessment of potential benefit from atezolizumab therapy.

#### Note 5: Final report generated by Medea

This example illustrates Medea's final report, which synthesizes outputs from the Analysis module, the LiteratureReasoning module, and the LLM backbone. Rather than relying on any single component (e.g., COMPASS's prediction), Medea reaches a consensus based on multiple evidence streams. By integrating immune concept interpretations derived from COMPASS, pathway analyses using Enrichr tools, and literature identified through context-verified retrieval, Medea uncovers immunological patterns that are inconsistent with responsiveness to atezolizumab.

Medea output - Final Analysis Report

**Response Likelihood to Atezolizumab in Bladder Urothelial Carcinoma**

**Assessment Based on Tumor Mutational Burden**

For the Caucasian female patient with bladder urothelial carcinoma and a tumor mutational burden (TMB) of 5.0 mutations/Mb, the estimated likelihood of a positive response to atezolizumab is approximately **15-20%**, with complete response rates likely below 10%.

**Scientific Rationale**

This estimate is based on the following key factors:

**1. TMB as a Biomarker:** The patient's TMB of 5.0 is considered low to intermediate. Clinical data from trials such as IMvigor130 indicate that TMB $>$ 10 mutations/Mb is associated with significantly better outcomes for atezolizumab monotherapy compared to chemotherapy. With a TMB of 5.0, which falls below this favorable threshold, response rates are expected to be more modest.

**2. Immune Microenvironment Analysis:** Transcriptomic analysis from the COMPASS tool indicated:

- Normal NK cell function (score: 0.3247)
- High T cell exhaustion (ratio: 1.61)
- Low B cell activity
- Moderate immune checkpoint score (0.7513)

These findings suggest an immune microenvironment that is not optimally primed for robust immunotherapy response.

**3. Sampling Limitations:** The transcriptomic analysis was performed on a kidney biopsy sample, which may not fully represent the bladder tumor microenvironment, potentially affecting prediction accuracy.

**Additional Considerations**

The prediction would be significantly refined with knowledge of:

- **PD-L1 Expression Status:** A critical determinant of atezolizumab response in urothelial carcinoma
- **FGFR3 Mutation Status:** FGFR3 mutations are associated with worse outcomes with immunotherapy in urothelial carcinoma
- **Molecular Subtype:** Luminal versus basal subtypes have different response patterns to immunotherapy
- **Prior Treatment History:** Response rates differ between first-line and later-line therapy

**Conclusion**

The estimated 15-20% likelihood of response is a preliminary assessment based primarily on COMPASS prediction results and TMB. For a more comprehensive prediction, additional biomarker testing would be recommended, particularly PD-L1 expression analysis, which could significantly alter this estimate in either direction.

#### Note 6: Evaluation of CellVoyager

For the benchmarking results on therapeutic target nomination, we evaluate the ability of CellVoyager [5] to nominate cell type specific therapeutic targets for rheumatoid arthritis (RA) by replicating the analytical procedures described in the PINNACLE study [6]. We evaluate CellVoyager across 420 analytical runs spanning seven distinct cell types as part of the Therapeutic Target Nomination benchmark. All experiment logs, execution metadata, and Jupyter notebooks containing CellVoyager's outputs are available on Figshare at: 10.6084/m9.figshare.30959984.

#### Note 7: Example abstention case from Medea

The following example illustrates an analysis by Medea (Claude 3.7 Sonnet) that results in an abstension due to weak statistical signals, such as negligible DepMap co-dependency scores (e.g., correlation coefficient of $-0.0335$ with $p$-value $= 0.223$), and the absence of shared biological pathways and direct empirical evidence in the literature. It illustrates Medea's ability to selectively abstain in uncertain cases, leading to more reliable synthetic lethality analyses.

User Instruction

We have introduced concurrent mutations in EME1 and RUNX1 in the MCF7 cell line. Could you clarify what synthetic lethal interaction might be observed in this context, if any?

Experiment Instruction

Use DepMap data to retrieve correlation metrics reflecting the co-dependency of the gene pair on cell viability. Next, perform pathway enrichment analysis with Enrichr to identify whether pathways associated with cell viability are significantly enriched and could be impacted by the gene pair. Synthesize the DepMap and Enrichr results, evaluate whether the combined perturbation of these genes is likely to induce a significant effect on cell viability, and find literature support if it exists.

Medea `(Claude 3.7 Sonnet) output`

**Synthetic Genetic Interactions Between EME1 and RUNX1 Mutations in MCF7 Cells**

Based on the available evidence, concurrent mutations in EME1 and RUNX1 in MCF7 cells may theoretically create synthetic genetic interactions, though direct empirical evidence for this specific combination is limited. The DepMap correlation coefficient between these genes is -0.0335, which is not statistically significant (p=0.223, adjusted p=0.415), suggesting no strong co-dependency pattern across cancer cell lines.

**Theoretical Basis for Potential Interaction**

The possibility of synthetic interaction stems from the complementary functional roles of these genes:

1. **EME1** is a key component of the MUS81-EME1 endonuclease complex essential for DNA repair, particularly in homologous recombination and resolution of stalled replication forks.

2. **RUNX1** functions as a transcription factor with established roles in transcriptional regulation and has been implicated in DNA damage response pathways.

**Potential Mechanisms of Synthetic Interaction**

If a synthetic interaction exists, it would likely operate through these mechanisms:

1. **Compromised DNA Damage Response**: The combined deficiency in EME1-mediated DNA repair and RUNX1-regulated transcriptional responses to DNA damage could create vulnerability where cells cannot adequately respond to endogenous DNA damage.

2. **Cell Cycle Checkpoint Dysfunction**: Both genes contribute to different aspects of cell cycle regulation, and their combined mutation may disrupt multiple checkpoint mechanisms.

3. **Increased Genomic Instability**: The combination could lead to elevated chromosomal aberrations beyond what either single mutation causes alone.

**Experimental Validation Approaches**

To confirm whether synthetic interactions exist between these mutations in MCF7 cells, consider:

- Assessing cell viability and proliferation rates in double mutants compared to single mutants and wild-type controls

- Measuring sensitivity to PARP inhibitors or DNA-damaging agents
- Quantifying unresolved DNA damage through $\gamma$H2AX foci analysis
- Examining chromosomal aberrations and micronuclei formation
- Analyzing cell cycle progression and checkpoint activation

The absence of shared pathways in the computational analysis suggests that any potential interaction may be based on complementary rather than overlapping functions. Therefore, experimental validation is essential before drawing definitive conclusions about synthetic interactions between EME1 and RUNX1 mutations in MCF7 cells.

#### Supplementary Tables

**Supplementary Table 1: Example user inputs from the three benchmark datasets for Therapeutic Target Nomination, Synthetic Lethality Prediction, and Immunotherapy Response Prediction.** Each analysis includes a user instruction ( grey ) and an optional experiment instruction ( green ) that specifies computational analyses to be performed by the agent. These examples illustrate the benchmarks' coverage of diverse reasoning tasks, including multi-choice, open-ended, and multimodal reasoning.

| Task Dataset | Sample |
|---|---|
| Therapeutic Target Nomination | Among POLD4, EDN3, RNU1-1, PAFAH1B3, and ASH2L, which gene exhibits the highest cell type specificity in naïve, thymus-derived $CD4^{+}$ $\alpha\beta$ T cells, making it the most promising candidate for targeting follicular lymphoma?<br>Identify and pull the verified follicular-lymphoma (FL) therapeutic targets and their PINNACLE embeddings; pull the PINNACLE activation gene set for naïve, thymus-derived $CD4^{+}$ $\alpha\beta$ T cells; intersect these sets to obtain the cell type specific FL targets. Calculate the average embedding for these overlapping targets and then compute the cosine similarity between this reference and each individual target gene. If one gene exhibits a markedly higher similarity than the others, nominate it as the primary candidate and provide a concise, evidence-based rationale for its biological relevance. |
| Synthetic Lethality Prediction | What would be the synthetic lethal interaction resulting from the concurrent mutation of RNF168 and APC in the MCF7 cell line, if any?<br>Use DepMap data to retrieve correlation metrics reflecting the co-dependency of the gene pair on cell viability. Next, perform pathway enrichment analysis with Enrichr to identify whether pathways associated with cell viability are significantly enriched and could be impacted by the gene pair. Synthesize the DepMap and Enrichr results, evaluate whether the combined perturbation of these genes is likely to induce a significant effect on cell viability, and find literature support if it exists. |
| Immunotherapy Response Prediction | I have mRNA TPM data characterizing the tumor immune microenvironment in a patient with Bladder Urothelial Carcinoma. What is the estimated likelihood of a positive response to Atezolizumab therapy? The patient's details are as follows:<br>• Sex: Female<br>• Race: Caucasian<br>• Tissue: Kidney<br>• Carcinoma: Bladder Urothelial Carcinoma<br>• Tumor Mutational Burden (FMOne mutation burden per MB): 5.0<br>Attachment: patient-62fb13-ar-62fb138.pkl<br>For patients with suspected immunodeficiency, begin by using the COMPASS tool. Start with an assessment of natural killer cell function, look for indicators of T cell exhaustion, and evaluate the activity of both B cells and plasma cells. Once you have calculated the COMPASS score, compare these results with transcriptomic markers of immune activation to identify those who may benefit from targeted therapies. |

**Supplementary Table 2: Accuracy (%) on the synthetic lethality prediction benchmark.** Each method is evaluated only on the subset of gene pairs for which it produces a definitive prediction. "Failed" predictions from agents are considered to be incorrect, and abstensions are excluded (refer to Supplementary Table 3 for abstention rates).

| Method | MCF7 | MDAMB231 | CAL27 | CAL33 | MCF10A | A427 | A549 |
|---|---|---|---|---|---|---|---|
| GPT-5.2 | 56.08 | 42.77 | 46.02 | 42.26 | 44.44 | 50.57 | 48.14 |
| PiLSL | 50.82 | 46.22 | 49.44 | 44.44 | 50.71 | 46.93 | 49.15 |
| Medea (GPT-5.2) | **62.03** | **64.15** | **56.77** | **51.46** | **52.81** | **57.54** | **57.20** |

**Supplementary Table 3: Abstention rate (%) on the synthetic lethality prediction benchmark.** The abstention rate measures the fraction of gene pairs on which the method declines to make a prediction, either by explicit abstention or by producing an uninterpretable output. PiLSL is a supervised classifier that always predicts a label (i.e., cannot abstain; denoted by "–").

| Method | MCF7 | MDAMB231 | CAL27 | CAL33 | MCF10A | A427 | A549 |
|---|---|---|---|---|---|---|---|
| GPT-5.2 | 19.13 | 31.08 | 37.22 | 33.61 | 28.21 | 30.69 | 18.71 |
| PiLSL | – | – | – | – | – | – | – |
| Medea (GPT-5.2) | 13.66 | 15.54 | 13.89 | 14.17 | 13.68 | 14.65 | 9.45 |

**Supplementary Table 4: Predicting synthetic lethality in *S. cerevisiae* under bleomycin and dimethyl sulfate conditions.** Left two columns report the model's accuracy on non-abstention predictions; right two columns report the model's abstention rate.

| | **Accuracy (%)** | | **Abstention Rate (%)** | |
|---|---|---|---|---|
| **Method** | Bleomycin | Dimethyl sulfate | Bleomycin | Dimethyl sulfate |
| GPT-5 | 33.33 | 39.09 | 9.30 | 10.23 |
| GPT-5.2 | 34.25 | 39.76 | 29.85 | 27.19 |
| Medea (GPT-5) | 48.26 | 51.74 | 22.09 | 15.79 |
| Medea (GPT-5.2) | **60.68** | **62.73** | 54.65 | 35.67 |

#### Supplementary Figures

Prompts for evaluating therapeutic target nomination

From the statement, extract the single strongest target gene being proposed/suggested. If a single gene is clearly recommended as top/primary/most promising, return that gene's symbol verbatim. If no unambiguous gene is proposed, output a label. Return only the target gene name or category label (no extra text).
Categories:

- Abstain: If evidence is insufficient/ambiguous, mixed without a clear winner, only non-gene entities (pathways/drugs/cell types) are proposed, or only gene families/classes are named without a specific gene.
- None: If the statement explicitly evaluates each candidate and disqualifies all of them for the context.
- The paragraph refuses, reports inability, or returns an error (e.g., "I can't help," "tool failed," "none" with no reasoning).

Statement: {reasoning_result}

**Supplementary Figure 1:** Prompt template for instructing the LLM judge to evaluate the nominated therapeutic targets.

Prompts for evaluating immunotherapy response prediction

Determine which category best describes the statement below. Return EXACTLY ONE category name (no extra text).
Categories:

- R: A positive verdict that response/benefit is likely (e.g., "likely/moderately likely to respond," "responder," "clinical benefit expected").
- NR: A negative verdict that response is unlikely (e.g., "unlikely to respond," "non-responder," "no benefit," "resistant").
- Abstain: Ambiguous or mixed evidence with no overall verdict; hedged assessments without a clear lean; requests more information.
- Failed: The paragraph refuses, reports inability, or returns an error (e.g., "I can't help," "failed," "none" with no reasoning).

Statement: {reasoning_result}

**Supplementary Figure 2:** Prompt template for instructing the LLM judge to evaluate immunotherapy response predictions.

Prompts for evaluating synthetic lethal prediction

Given a statement, determine which category below it fits, based on whether it concludes that synthetic lethality (SL) represents a (potential) genetic interaction between two genes. Return EXACTLY ONE category name (no extra text).
Categories:

- Synthetic lethality: The paragraph clearly concludes or strongly supports that the combined perturbation (loss/inhibition/KO) of the two genes reduces viability, and concludes that synthetic lethality (SL). Signals include: "synthetic lethal/sickness," "double knockout is lethal," "co-inhibition is lethal/toxic"
- Non-SL: The paragraph clearly concludes that synthetic lethality (SL) is unlikely.
- Abstain: Ambiguous, mixed, or insufficient evidence; hedged without a clear verdict; mentions pathways/contexts but no firm conclusion.
- Failed: The paragraph refuses, reports inability, or returns an error (e.g., "I can't help," "failed," "none" with no reasoning).

Statement: {reasoning_result}

**Supplementary Figure 3:** Prompt template for instructing the LLM judge to evaluate synthetic lethality predictions.

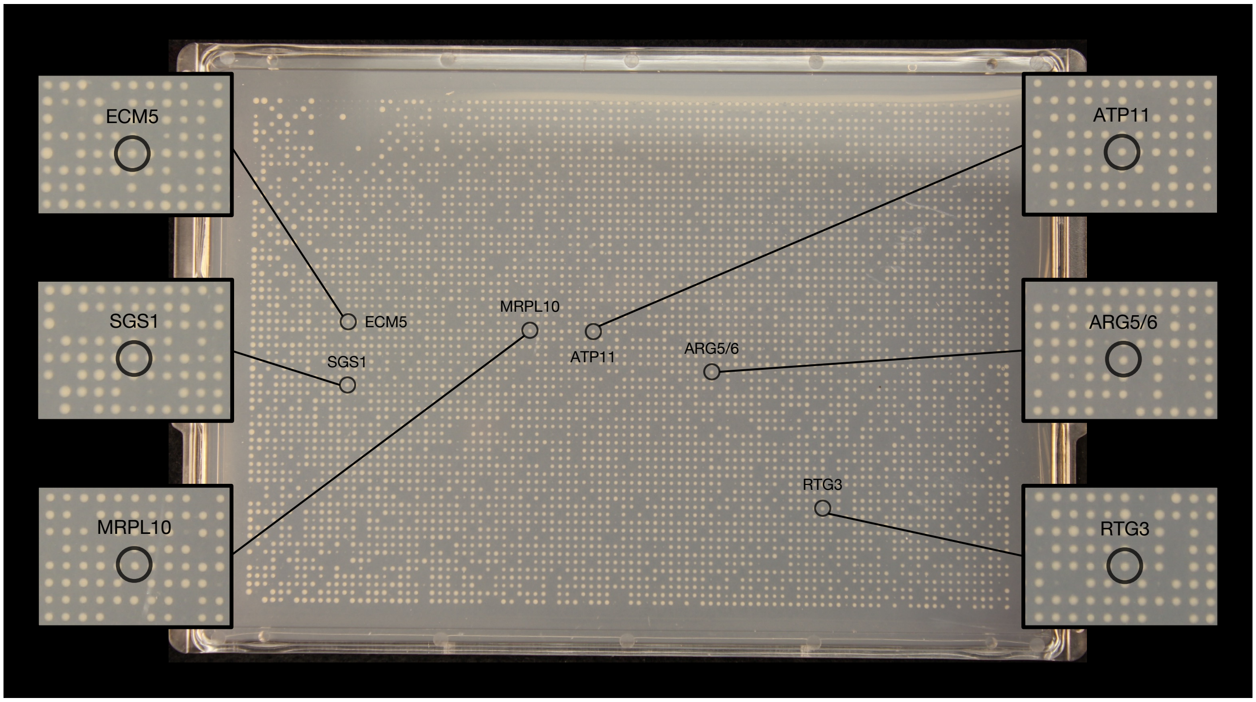


**Supplementary Figure 4: Single knockouts of SGS1 interaction partners are viable under dimethyl sulfate.** A 6,144-colony plate from an Epistatic MiniArray Profile screen profiling crosses between the genome-wide library of yeast single-deletion strains and a query strain deleted for HO, a dispensable endonuclease locus used as a neutral control, grown under dimethyl sulfate. Because HO deletion carries no fitness cost, colony size at each array position reports the fitness of that single knockout alone under this condition. Circles mark the array positions of SGS1 and its five predicted interaction partners from Figure 6d: MRPL10, ARG5/6, ECM5, RTG3, and ATP11. All six form viable colonies, confirming that the loss of viability observed for the corresponding double knockouts with SGS1 (Figure 6d) reflects the combined perturbation rather than a single-gene fitness defect under dimethyl sulfate.
